# Tree segmentation in airborne laser scanning data is only accurate for canopy trees

**DOI:** 10.1101/2022.11.29.518407

**Authors:** Yujie Cao, James G. C. Ball, David A. Coomes, Leon Steinmeier, Nikolai Knapp, Phil Wilkes, Mathias Disney, Kim Calders, Andrew Burt, Yi Lin, Tobias D. Jackson

## Abstract

Individual tree segmentation from airborne laser scanning data is a longstanding and important challenge in forest remote sensing. There are a number of segmentation algorithms but robust intercomparison studies are rare due to the difficulty of obtaining reliable reference data. Here we provide a benchmark data set for temperate and tropical broadleaf forests generated from labelled terrestrial laser scanning data. We compare the performance of four widely used tree segmentation algorithms against this benchmark data set. All algorithms achieved reasonable accuracy for the canopy trees, but very low accuracy for the understory trees. The point cloud based algorithm AMS3D (Adaptive Mean Shift 3D) had the highest overall accuracy, closely followed by the 2D raster based region growing algorithm Dalponte2016+. This result was consistent across both forest types. This study emphasises the need to assess tree segmentation algorithms directly using benchmark data. We provide the first openly available benchmark data set for tropical forests and we hope future studies will extend this work to other regions.

## 1 Introduction

Airborne laser scanning (ALS) is widely used in forest ecology, but automatic individual tree segmentation (ITS) in dense broadleaf forests remains challenging (1). Accurate ITS algorithms would enable researchers to study tree growth (2), mortality (3), leaf phenology (4) and carbon dynamics (5) remotely, providing opportunities to track changes at landscape scales. These complement existing field-based methods, which are essential but limited in scale (6). Many ITS algorithms have been developed, but robust comparisons of their accuracy are lacking, particularly for broadleaf forests. In this study we generate an ALS tree segmentation benchmark data set and use it to assess leading ITS algorithms for temperate and tropical broadleaf forests.

There are two broad categories of ITS algorithms: 2D raster algorithms and 3D point cloud algorithms. Raster ITS algorithms are based on 2D top of canopy height rasters, meaning that information about understory trees is excluded and subcanopy trees cannot be segmented. A widely used 2D-raster ITS algorithms is the region growing algorithm of Dalponte et al. 2016 (7) (hereforth *Dalponte2016*). which has been applied across many forest types (8–11). Other 2D-raster ITS algorithms use clustering or image object detection methods like K-means (12), Pouring algorithm (13), Centroidal Voronoi Tessellation (14) and other variants (15–17).

Recent developments have focused on analysing the 3D point cloud ITS algorithms searching for clusters of points which may represent tree crowns including those of understory trees. These methods include rule-based and data-driven methods. Rule-based methods extract individual trees with a series of user-defined spatial constraints such as relative distances between trees, shape indices calculated from horizontal projections and point density changes (18–20). However, these methods often over-simplify forest structure and are thus not applicable to diverse forest types. Data-driven approaches rely on unsupervised clustering algorithms to extract tree boundaries. They include prior knowledge of forest structure before (21–24) or after (25) the clustering stage for the purpose of initialization or crown refinement.

Computational cost is an important consideration when working with large datasets (billions of points). For example, AMS3D has been widely used in multiple forest contexts, but it takes much longer running time compared with the 2D-raster counterparts like Dalponte2016. Potential solutions could be voxelization and down-sampling strategy before performing segmentation (18), but these could also lead to information loss.

Previous efforts to assess ITS algorithms have been limited by the available ground truth data. Many ITS algorithms were assessed based on their ability to predict some property of the forest such as: the number of trees (26, 27), tree trunk position (28),tree height (29), crown size (30, 31), diameter at breast (7, 32), stem volume (17) or above ground biomass (7, 33). However, ITS algorithms tuned to predict these forest properties may actually segment individual trees with low accuracy. In addition, ITS algorithms tuned in this indirect way are unlikely to generalize well and may produce unexpected results when applied to a new data set.

Direct ITS algorithm assessment studies have used manually interpreted point clouds (23) and segmented tree crowns (34), which allows the algorithms to be assessed directly on their ability to segment trees. However, manually segmented tree crowns are usually biased towards the large visible, canopy trees only (35). To the best of our knowledge all available benchmark data sets focus on conifers or temperate deciduous with relatively open canopy (36, 37). Virtual laser scanning provides highly detailed simulated ALS data, and has been used to assess ITS algorithms (38, 39). However, assessment on real data (i.e. not simulated data) is still required to evaluate the performance of ITS algorithms (40) in practical terms.

In this study, our key objectives are to:

1. Generate an ALS benchmark dataset with individual trees labelled in both the canopy and understory and make this openly available online.
2. Use this benchmark data to compare the performance of ITS algorithms.
3. Explore the sensitivity of each ITS algorithm to its key input parameters.

## 2 Materials and methods

### 2.1 Field Sites

Temperate broadleaf forests (Wytham Woods, UK) and Tropical rainforest (Sepilok Reserve, Malaysia), were included in this study (see Fig. 1.).

**Fig. 1.**
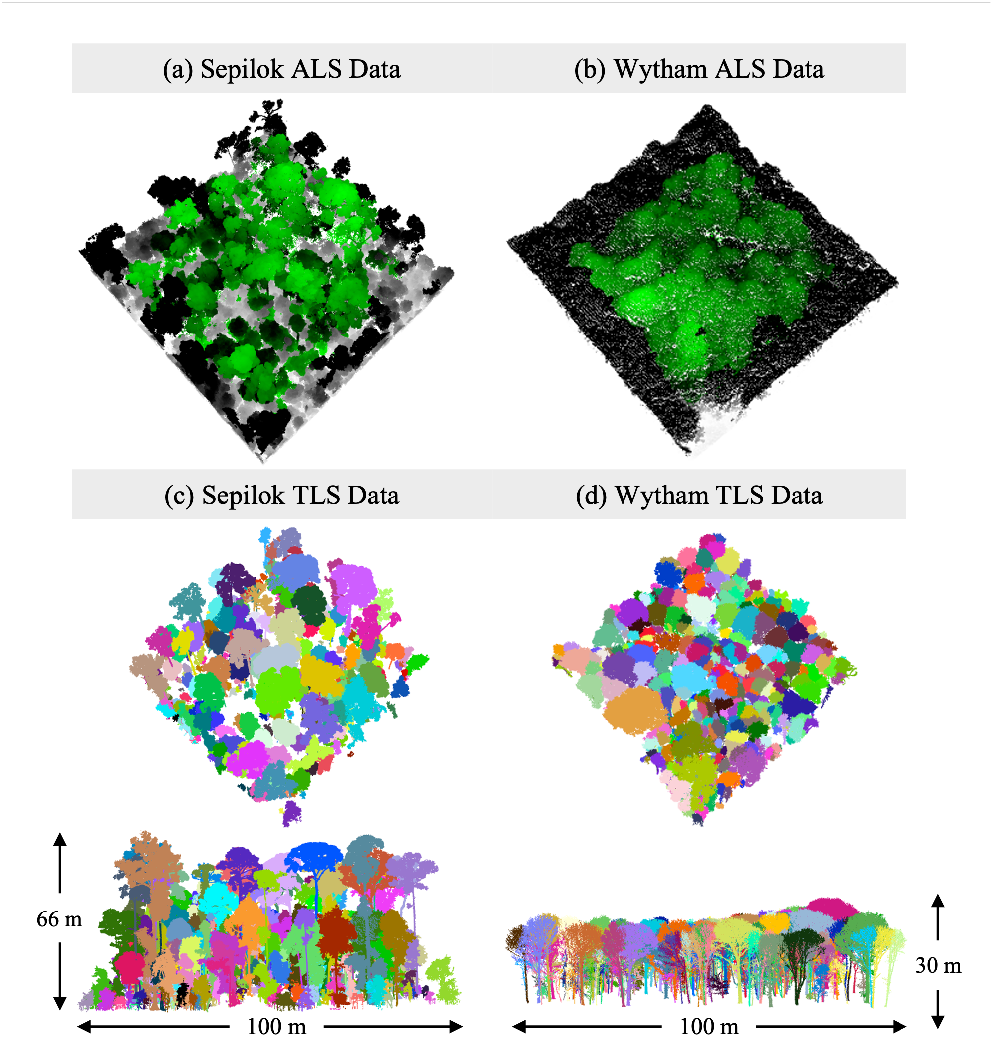
Raw ALS data and segmented TLS data of Sepilok Forest (left panel) and Wytham Woods (right panel): Green areas in (a) and (b) are core areas of raw ALS datasets; Black areas in (a) and (b) areas are buffered to obtain complete ALS individual tree point clouds for further algorithm assessment; (c) and (d) are labelled TLS datasets, both of which were used for creating ALS benchmark datasets.

Wytham Woods is a semi-natural woodland, located on a gentle hill with the maximum elevation rising to 165m in Oxfordshire, England (1° 20’ W, 51° 47’ N) (41). Since 2008, an 18 ha permanent inventory plot was established, which was the focus of this study(42). The woodland is mixed deciduous forest, dominated by Acer pseudoplatanus (Sycamore), Fraxinus excelsior (Ash), Corylus avellana (Hazel) and Quercus robur (Oak). The trees are diverse in terms of size and distribution with a maximum diameter at breast height of 141.2 cm, and the tallest tree is about 40 m. The 1ha plot involved in our study is a subplot with the maximum height reaches around 30 m.

Sepilok Forest Reserve (117° 56’ E, 5° 10’ N) is a remnant of lowland tropical rain forest located close to the north-east coast of Sabah, Malaysia (43). It is one of the oldest protected tropical forest in Southeast Asia, covering around 4500 ha with lowest and highest elevations of 50 m and 250 m respectively (44). The 1 ha plot involved in our study is nested in alluvial dipterocarp forest where over 160 species have been found. The maximum diameter at breast height is 159.2cm, while the tallest tree is around 66 m. (43)

### 2.2 LiDAR data

#### ALS data

In Sepilok, ALS data was acquired in February 2020. For the focal area, the ALS data has a pulse density of about 139 per m^2^. In Wytham Woods, ALS data was collected in summer 2014. Over the area of interest, it has a pulse density of around 6.26 per square metre (see table 1).

**Table 1.** TLS and ALS data details for Sepilok Forest and Wytham Woods

|  | Sepilok Forest | Wytham Woods |
| --- | --- | --- |
| Plot Area (ha) | 1 | 1 |
| Number of trees | 430 | 523 |
| TLS point density(pts/m<sup>3</sup>) | 5400 | 99 |
| ALS point density(pts/m<sup>3</sup>) | 139 | 7 |
| ALS-TLS alignment RMSE (m) | 0.192 | 0.625 |

#### TLS data

At both locations TLS data was captured using a RIEGL VZ-400 (RIEGL, Horn, Austra) following standard protocols described in (45). The instrument has a beam divergence of 0.35 mrad and operates in the infrared (wavelength 1550 nm) with a range up to 350 m, a pulse repetition rate of 300 kHz and the angular sampling resolution was 0.04°. At each scan position, two scans were acquired where the scanner rotation axis was perpendicular and parallel to the ground surface respectively. In both sites, reflective targets were located between scan positions to aid with coregistration (45). Scans were registered in RiSCAN Pro in a two-step process where (1) the reflective targets were used to generate a coarse registration (2) a set of planes generated from the point cloud were used in a Multi-station adjustment to improve the registration. The point cloud was then downsampled using a voxel size of 0.026 m and 0.02 m for Wytham and Sepilok respectively.

At Wytham, scans were done within a larger 6 ha area to ensure the best possible data quality within our 1 ha study area (Calders et al. 2022). TLS data were collected throughout December 2015 and January 2016 (in leaf-off conditions) on a 20 m x 20 m grid (46). Individual trees were segmented from the larger point cloud using *treeseg* (47) and manually checked for quality assurance.

In Sepilok, the study area was located in the tall alluvial forest and the data were collected in 2017. The local 1 ha plot reference is SP292/1 and SEP12 in the forestplots.net data base. TLS data for was captured across a 1 ha plot from 121 scan positions on a 10 m x 10 m grid. Trees were segmented from the point cloud using TLS2trees (Wilkes et al. in prep). Briefly, trees are segmented using a 2 step process where a semantic segmentation is used to classify points into ground, leaf, wood and coarse woody debris. Using the wood and leaf classes only, a graph based instance segmentation is used where woody stems are first identified, leaf points were subsequently added to individual stems. Resulting segmented trees were manually checked for quality assurance.

### 2.3 Creating ALS benchmark data sets

We aligned the ALS and TLS data using a geo-coordinate transformation with 9 manually selected feature points. These feature points were selected as the intersection of two branches in the upper canopy, so they could be easily identified in both ALS and TLS. The root mean square error in alignment for Sepilok was (0.192 m) and for Wytham was (0.625 m). We created a polygon to outline the core area in which the TLS (1ha) overlaps with the ALS (> 100 ha). To enable fast 3D data retrieval we used a *kdTree* (48), which is a space-partitioning data structure for organizing points in a *k* -dimensional space.

We labelled the individual trees in the ALS data using the segmented TLS data in order to produce a benchmark ALS data. Here, we used an iterative nearest neighbour voting strategy to label the ALS data. The labelling strategy consisted of the following steps (see Fig.2):

**Fig. 2.**
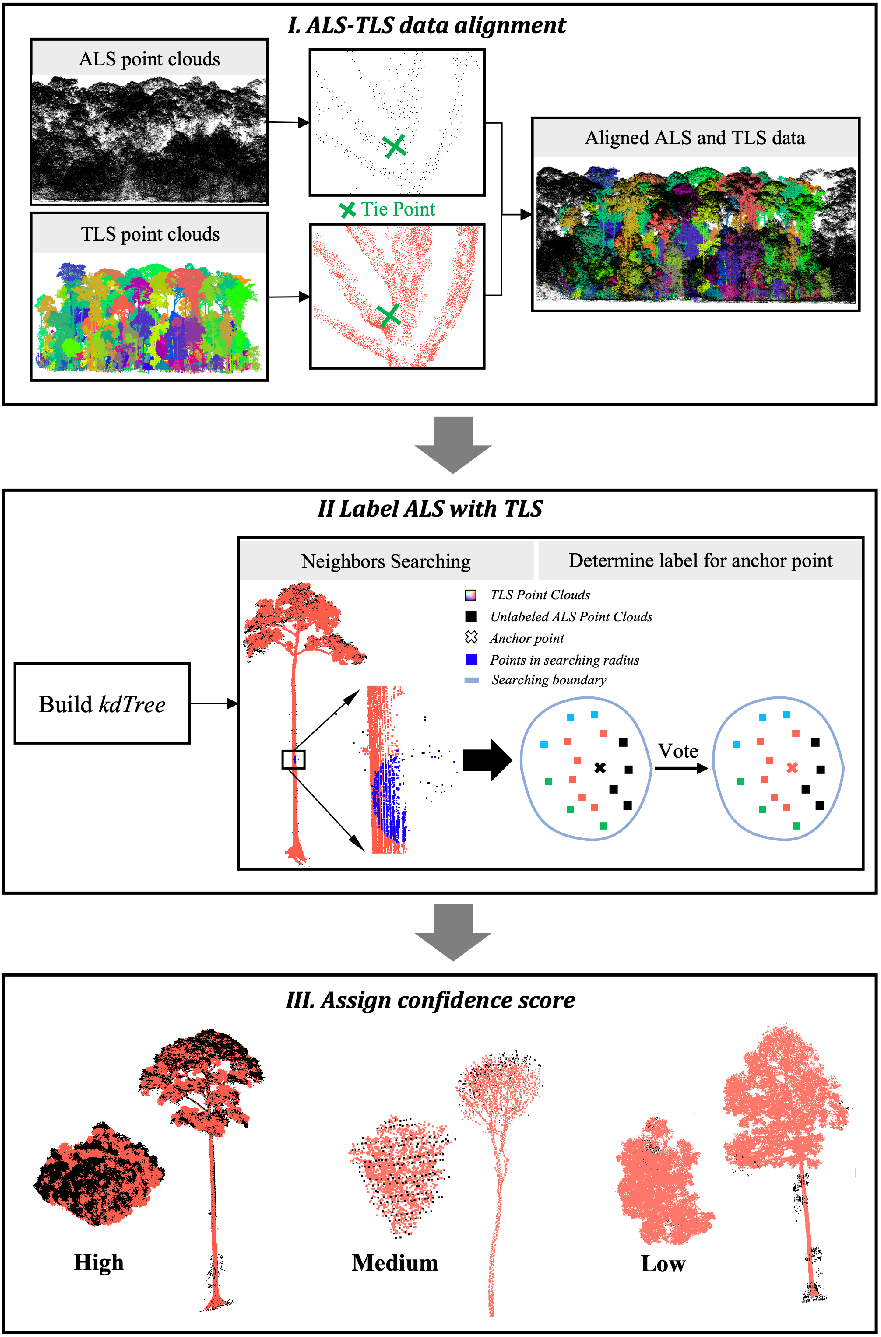
Workflow to create ALS benchmark with detailed TLS data: ALS and TLS point clouds were firstly aligned with manually selected feature point pairs; then a kdTree was built on ALS and TLS data space to speed up the retrieving process; ALS point clouds were labelled by a neighbour voting process and ALS benchmark datasets were then obtained where colorized areas are core areas and black areas are buffered areas; After that, individual tree point clouds from the ALS benchmark were given one of the seven confidence scores (see Supplementary B) by manually checking the data quality.

1. For each ALS point, we selected a maximum of 900 (Sepilok) or 300 (Wytham) surrounding points within a radius of 1.4 m (Sepilok) or 2.0 m (Wytham).
2. The label of each ALS point was then assigned as the label of majority of these neighbour points.

This process was repeated until it converged (i.e. the number of newly labelled ALS points was less than 5). The pseudocode demo for ALS labelling strategy can be found in Algorithm 1.

### 2.4 Manually rating ALS benchmark trees with confidence score

A confidence score system was introduced to control the data quality of the labelled ALS benchmark tree (see Fig.2 as well as Supplementary C and D). We categorized the trees as high, medium and low confidence. The scores were defined in terms of the number of points contributing to a crown polygon, the distribution of ALS point clouds with comparison to corresponding TLS data and bias of individual tree feature representation. All points within a tree were assigned the tree-level confidence score. For a detailed description of the confidence score system see supplementary B.

#### Algorithm 1 ALS Labelling Strategy

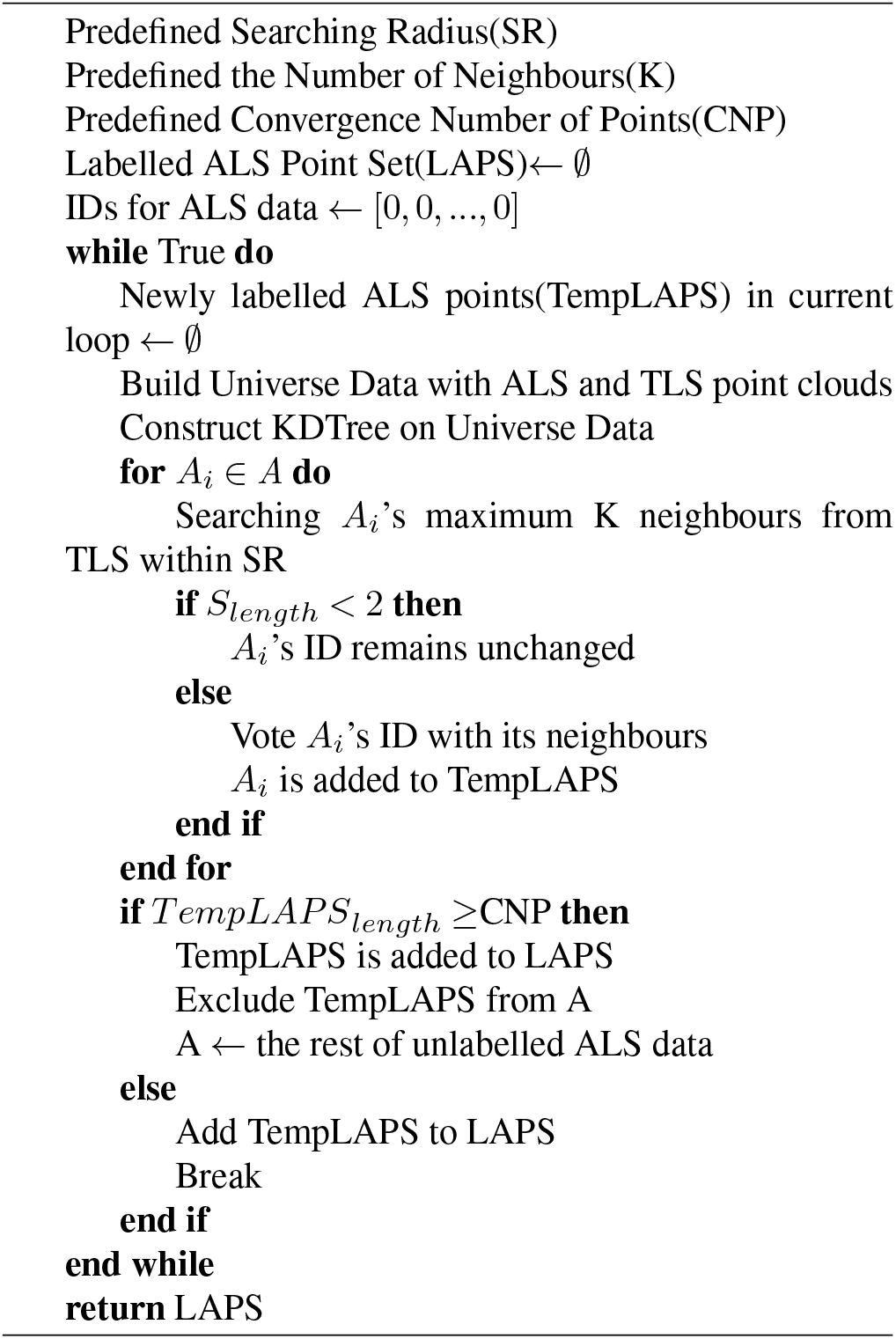

In Sepilok, we noticed four large trees which were clearly visible in the ALS data, but not in the TLS. This is presumably because the trees had fallen in the 3 years between the scans, and these trees were assigned a low confidence score.

Importantly, ITS algorithms segmented trees on the raw ALS data, but only trees belonging to high and medium classes were used in our assessment. The confidence score of the predicted trees was assigned as the most common confidence score of the points within it. We filtered by confidence score during our assessment process to ensure the results are not due to poor quality reference data. Finally, the ALS benchmark for Sepilok Forest and Wytham Woods were created (see Fig. 3)

**Fig. 3.**
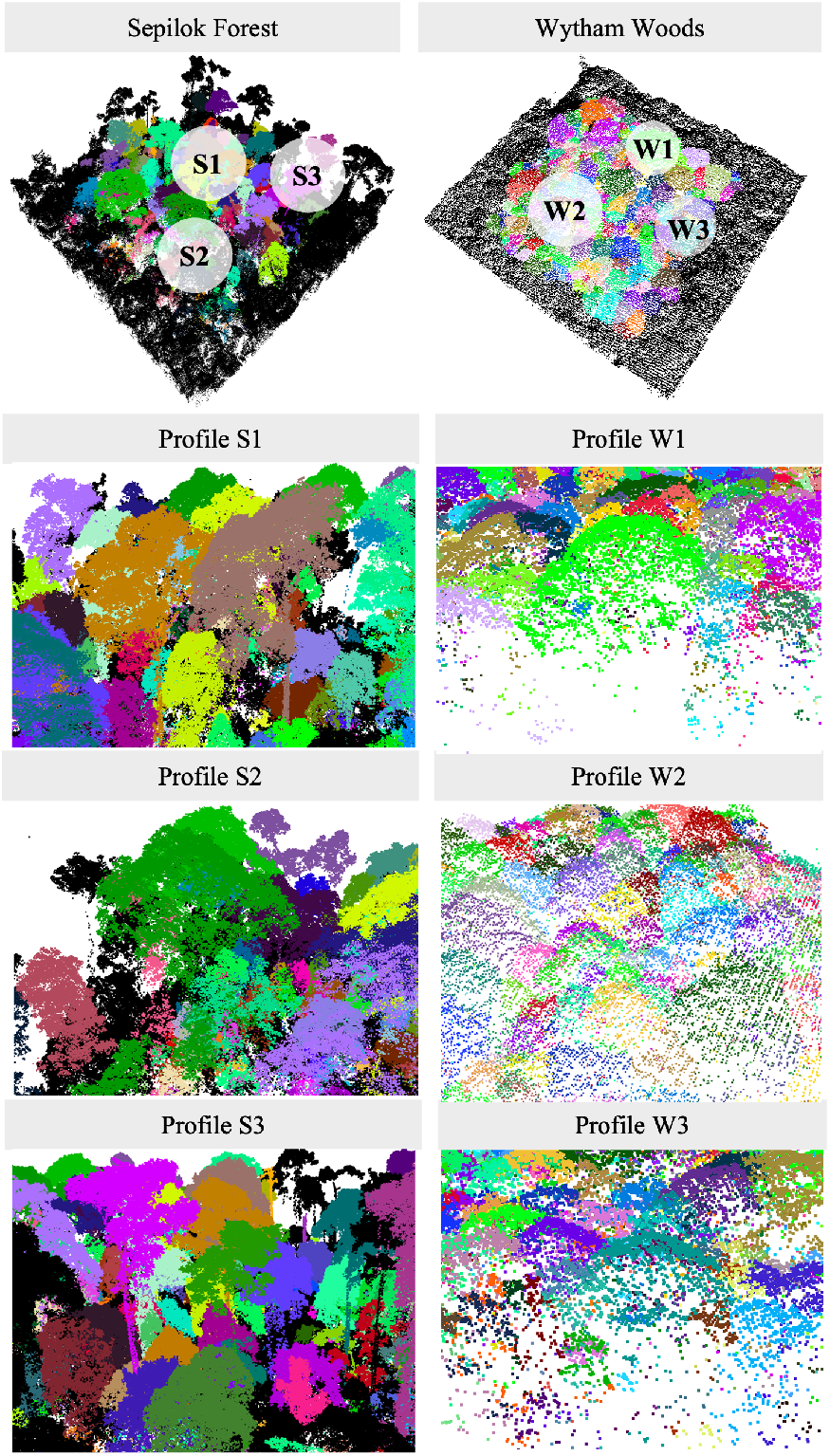
ALS benchmark data: Left panel displays ALS data for Sepilok Forest, and Profile S1, S2 and S3 show details of the data; Right panel is ALS data for Wytham Woods; Profile W1, W2 and W3 present details of the data.

### 2.5 ITS algorithm selection

We reviewed all highly cited ITS algorithms and summarized our findings in Table 2. We noted the algorithm type (2D or 3D), key methods, assessment method, forest type and overall accuracy and the number of citations. We chose to focus on four of the most highly cited algorithms, which are representative of different algorithm structures.

**Table 2.** Review of 10 highly cited ITS algorithms: DBH is diameter at breast height, while ACD is aboveground carbon density. Where multiple accuracy metrics are given we report the best one for each algorithm. A more detailed ITS algorithm review can be found in Supplementary A

| Author/Method Name (year) | Citations 2020-22 | Algorithm Type | Forest Type | Reference Data | Trees | Main Result |
| --- | --- | --- | --- | --- | --- | --- |
| Dalponte2016 (7) | 108 | Raster | Temperate broadleaves | Field measured DBH | 1762 | RMSE: 11 cm |
| Dalponte2016+ (52) | 76 | Raster | Tropical broadleaves | Field measured ACD | 91 | RMSE: 13% |
| Li2012 (19) | 219 | Point | Conifers | Field measured tree location | 380 | F1 score: [0.83, 0.95] |
| AMS3D (21) | 88 | Point | Topical broadleaves | Field measured aboveground biomass | 1604 | RMSE:[4.2%, 16.9%] |
| Chen2006 (16) | 132 | Raster | Temperate broadleaf | Manually delineated tree crown | 772 | Absolute Accuracy: [56%, 61%] |
| Silva2016 (14) | 104 | Raster | Conifers | Field measured basal area | 185 | RMSD: 0.07 |
| Hyypä2001 (17) | 99 | Raster | Conifers | Field measured basal area | 15 | System Error: +3.9m2/ha |
| Dai2018 (23) | 63 | Point | Temperate broadleaf | Manually labelled point clouds | 392 | Average Accuracy: 84% |
| LayerStacking2017 (22) | 57 | Point | Temperate broadleaf | Field measured tree location | 1962 | Commission Error: [22%, 53%] |
| Williams2019 (32) | 28 | Point | Tropical broadleaves | Field measured aboveground biomass | 91 | RMSE:[14%, 22%] |

### Dalponte2016 and Dalponte2016+

are 2D-raster based ITS algorithms. Dalponte2016 is widely used as it is integrated in the lidR R package (7, 49). The algorithm firstly finds treetops with local maxima filtering on a top of canopy height raster and determines the tree crown boundary with a region growing and decision tree method. In detail, a pixel is viewed as a treetop if its Z value is higher than its surroundings inside the sliding windows. Then the crown will grow around the treetop. During the region growing, TH_CR and TH_SEED are key parameters contributing to the final tree crown polygon. In detail, TH_SEED is factor controlling the height difference between the treetop and neighbouring pixels. A neighbouring pixel would be included in the region if its height is greater than the product of TH_SEED and the height of the treetop. While, TH_CR controls the difference between the mean height of the regions and their surrounding pixels, and a neighbouring pixel would be view as part of the region if its height is greater than the product of TH_CR and the mean height of the growing region (see table 3). Therefore, the range for the two parameter ranges from 0 to 1. The code for Dalponte2016 used in the study can be found in (50, 51).

**Table 3.**
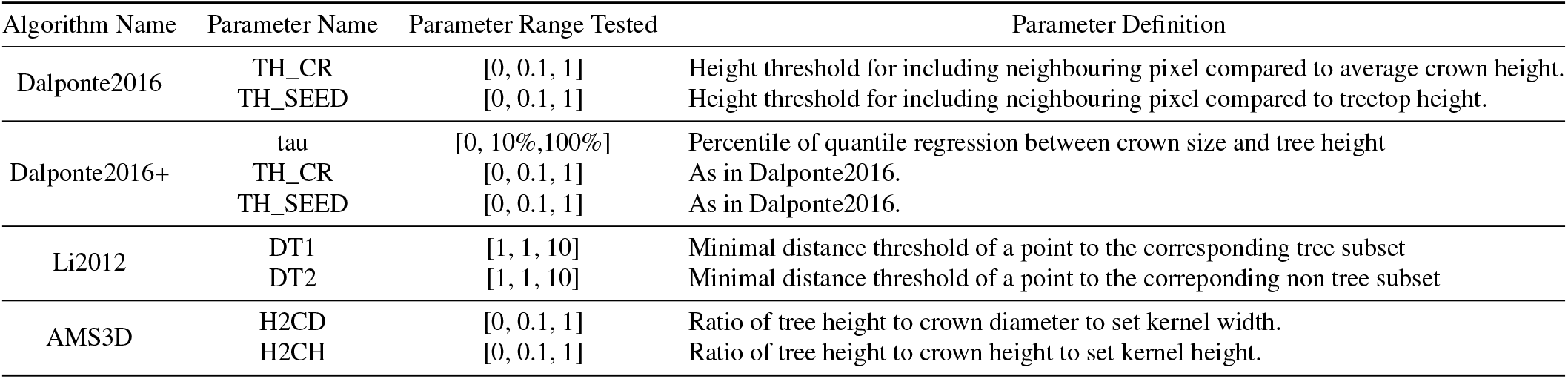
Details for ITS algorithms and the parameters assessed in the experiment.

| Algorithm Name | Parameter Name | Parameter Range Tested | Parameter Definition |
| --- | --- | --- | --- |
| Dalponte2016 | TH_CR | [0, 0.1, 1] | Height threshold for including neighbouring pixel compared to average crown height. |
|  | TH_SEED | [0, 0.1, 1] | Height threshold for including neighbouring pixel compared to treetop height. |
| Dalponte2016+ | tau | [0, 10%, 100%] | Percentile of quantile regression between crown size and tree height |
|  | TH_CR | [0, 0.1, 1] | As in Dalponte2016. |
|  | TH_SEED | [0, 0.1, 1] | As in Dalponte2016. |
| Li2012 | DT1 | [1, 1, 10] | Minimal distance threshold of a point to the corresponding tree subset |
|  | DT2 | [1, 1, 10] | Minimal distance threshold of a point to the corresponding non tree subset |
| AMS3D | H2CD | [0, 0.1, 1] | Ratio of tree height to crown diameter to set kernel width. |
|  | H2CH | [0, 0.1, 1] | Ratio of tree height to crown height to set kernel height. |

By introducing a flexible searching window, the improved Dalponte2016 (herein *Dalponte2016+*) (52) is able to find more realistic treetops before growing stage. The variable window changes according to the crown size and tree height relation. The crown size tends to be smaller for shorter trees and thus the searching window is smaller. In order to find the optimal the crown size and tree height relation for different forest structures, the global allometry database (53) was used. In the database, Wytham Woods and Sepilok Forest belong to Palearctic and Indomalayan realms respectively. The crown size and tree height relations can be described by quantile regression on allometry of different biogeographic realms. The parameter tau means different percentile of quantile regression, which represents different crown size and tree height relations. As for TH_SEED and TH_CR, they share the same definition with those in Dalponte2016.

**Li2012** (19) is a rule-based point ITS algorithm, which has been integrated in the LiDAR processing packages (51) and commercial software (54). This ITS algorithm starts with finding the global highest point in the data space. Then the highest point and a dummy point far away from the highest point are used to create two sets. The rest of points are assigned to the two sets following a top-down order, relative spacing, shape index and point density distribution. In the procedure, DT1 and DT2 are the key parameters (see table 3), which present the minimal distance to tree set and non tree set respectively. The range of the two parameters is from 1 to 10. Point clouds of individual trees can be obtained by repeating above procedure till no unlabelled point left. The code for Li2012 used in our experiment can be found in LidR package(49).

### Adaptive Meanshift 3D (AMS3D)

was implemented according to the descriptions by (21, 55). It is based on the general principle of mean shift clustering (56), applied to the 3-dimensional coordinate space of the lidar point cloud and adapted to the fact that tree crowns in upper canopy strata are larger than in lower strata. For every point in the point cloud the center of point density in a cylindrical search neighborhood (the kernel) is identified. The weight with which each neighbor point contributes to the calculation of the center depends on its relative position inside the kernel. The kernel is then shifted to this center of point density. This procedure is iteratively applied until the kernel reaches a stable position, which usually is closely underneath the apex of a tree crown. After a final kernel position for every point in the point cloud has been found, tree crown clusters are formed from all points for which the kernel positions converge at the same crown apex. Since the final kernel positions do not fully converge, the DBSCAN clustering algorithm (57) was applied to the final kernel positions to identify clusters. To enable fast AMS3D computations, the algorithm was implemented in C++ using an R*-tree spatial index structure for efficient spatial queries (58)(Leon MA citation).

The two parameters of the algorithm control the size of the kernel (cylinder height and diameter) as a function of kernel center height above ground. This height dependence ensures large crown clusters in the upper and small crown clusters in the lower canopy. Thus, typical crown length and crown diameter to tree height ratios are good choices for the parameters.

### 2.6 Assessing ITS algorithm accuracy against benchmark

#### Grid search over all parameters

We implemented each algorithm multiple times varying the key parameters using a grid search method to give a comprehensive overview of the sensitivity to these parameters, see table 3. The outputs were assessed by how well they matched tree crown polygons from the benchmark data set (see Fig.4).

**Fig. 4.**
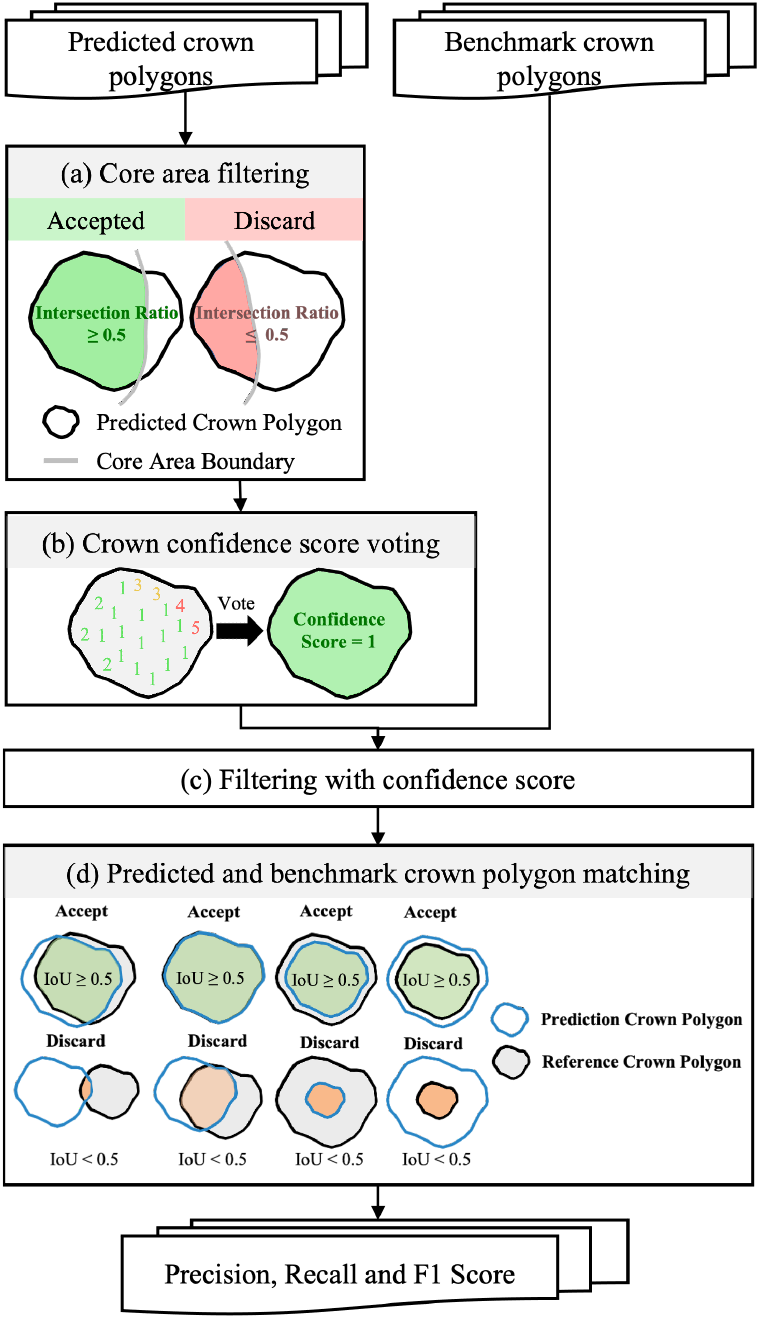
Crown polygon-based assessment framework for parameter tuning and method inter-comparison. (a) a filtering was conducted to select the predicted tree crowns with over 0.5 intersection rate with the core area; (b) A confidence score voting was then conducted to decide the confidence score for these predicted tree crown polygons; (c) Only those polygons with high or medium confidence score were used for assessment; (d) During tree crown polygon matching, the predicted polygons would be accepted as TP if its IoU with the corresponding benchmark crown polygon was larger than 0.5.

#### Assessment workflow

The assessment was carried out using polygons to represent the individual tree crowns from both the benchmark and predicted trees (59, 60). During assessment (see Fig.4), predicted tree crown polygons with more than half its area inside the core plot area (the overlap between TLS and ALS) were selected as candidates for further assessment. A voting stage was then implemented to decide the confidence score for a predicted tree crown as different confidence scores could be included in a single predicted tree crown polygon. After that, the input for polygon matching could be selected with targeted confidence combinations. During polygon matching, the IoU of a predicted tree crown polygon was determined by selecting the maximum IoU with corresponding reference tree crown polygon. A predicted tree crown polygon could be accepted as true positive if its maximum IoU ≥ 0.5. Precision, Recall and F1 Score could be finally obtained through true positive, false positive and false negative, all of which will be introduced in detail in the following sections.

These trees and all others with low confidence scores were retained during segmentation but filtered out prior to the assessment stage.

#### Assessment index

Accuracy metrics used in the study are based on IoU value of individual tree crown polygons, which is calculated as intersection over union (Equation 1). The predicted crown polygons with ≥0.5 IoU (Equation 2) were viewed as true positive (TP) (Equation 3). Those predictions failed to match a benchmark crown polygons stand for false positive (FP), while those benchmarkd tree crowns with no corresponding crown polygons in the prediction were classified into false negative (FN). Finally, TP, FP, FN were used for obtaining Precision (Equation 4), Recall (Equation 5) and F1 Score (Equation 6).

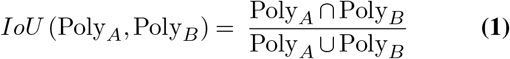

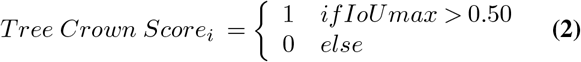

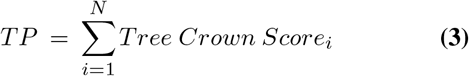

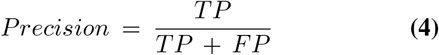

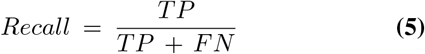

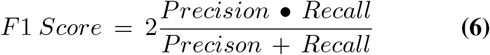

## 3 Results

### 3.1 Novel benchmark data set

We produced a novel benchmark data set for ITS algorithm assessment using TLS to label individual trees in ALS data (Figure 2). We manually assigned confidence scores to each tree in the benchmark data set. 170 trees in Sepilok and 139 in Wytham had a ‘high’ confidence score. Many canopy trees showed a visually perfect match between the TLS and ALS labels and taller trees generally higher confidence scores, mainly due to better ALS coverage. Importantly, our data set also contains many understory trees with ‘high’ or ‘medium’ confidence scores (75 trees in Sepilok Forests and 11 in Wytham Woods, see Figure B.1 in Supplementary B). This enables us to assess ITS algorithm performance for understory trees.

We tested the sensitivity of our results to the inclusion of trees with different confidence scores and found that our results were robust. Specifically, we found little change in results when only including trees with the highest confidence scores, compared to additionally including trees with medium confidence scores. The trees with low confidence score were considered too poorly matched between the ALS and TLS to be used for assessment and we found our results were sensitive to the inclusion of these trees. The complete matrix plots for Precision, Recall and F1 Score of the ITS algorithms can be found in Supplementary E and F.

### 3.2 Segmentation accuracy improves with tree height

ITS algorithm accuracy increased with tree heights across both sites (see Fig.5 and Fig.6).

**Fig. 5.**
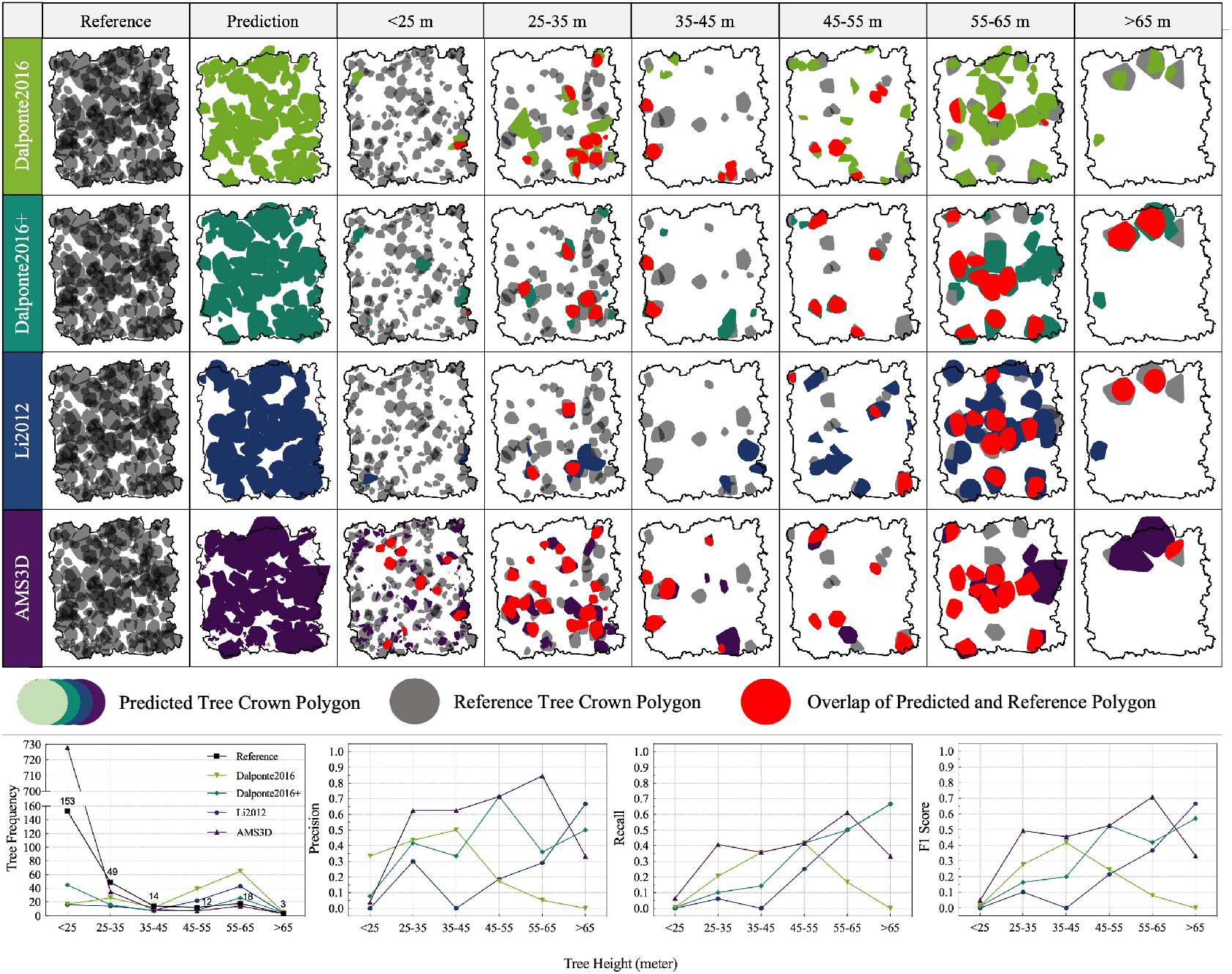
Algorithm performance changes with regards to tree heights in Sepilok Forest. The upper panel visualizes ALS benchmark polygons (grey), predicted polygons from 4 involved algorithms (vridis scale color) and intersection polygon; The buttom panel displays the statistical line-chart for tree frequency, precision, recall and F1 score.

**Fig. 6.**
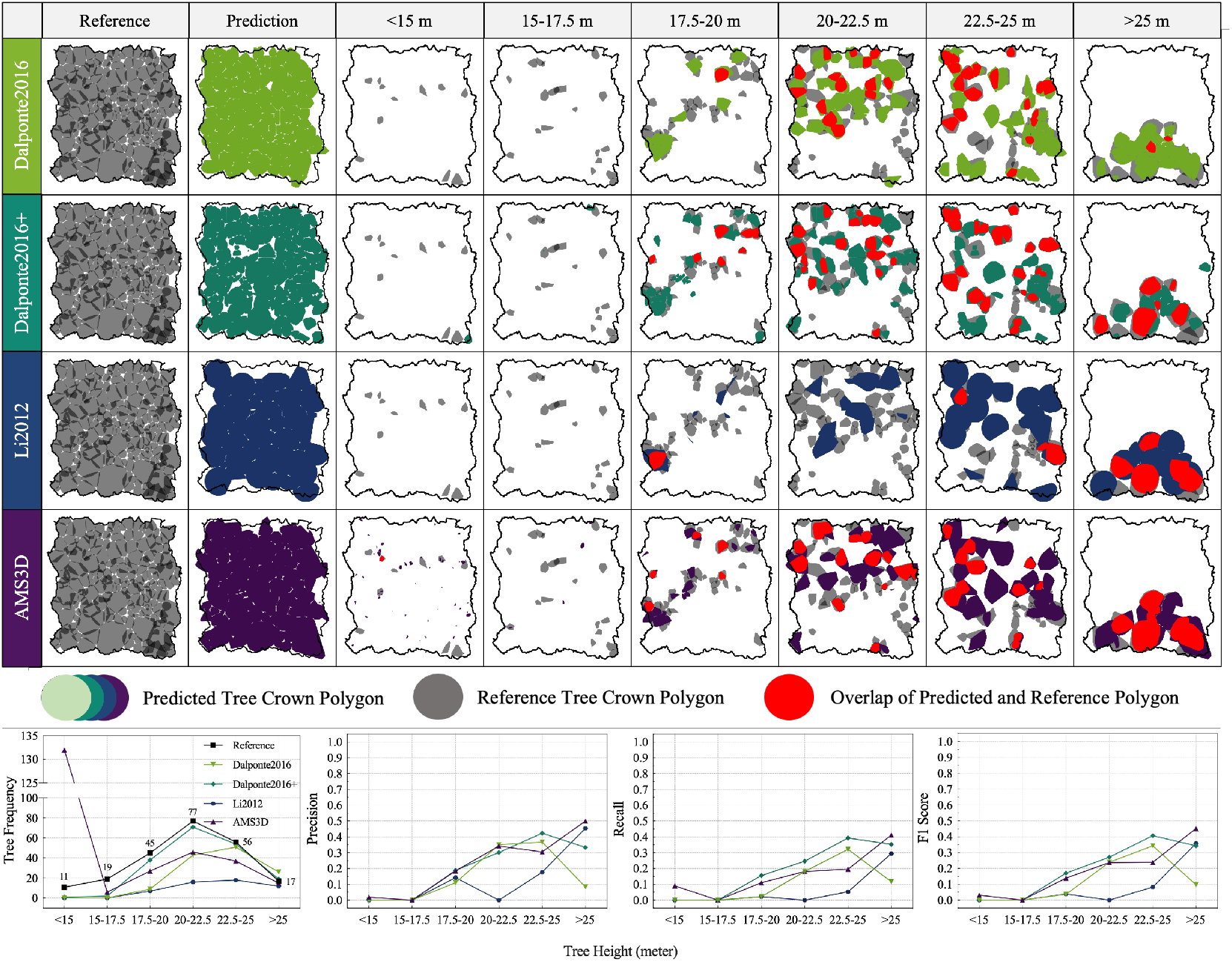
Algorithm performance changes with regards to tree heights in Wytham Woods. The upper panel visualizes ALS benchmark polygons (grey), predicted polygons from 4 involved algorithms (vridis scale color) and intersection polygon; The buttom panel displays the statistical line-chart for tree frequency, precision, recall and F1 score.

In Wytham, the precision and recall of all four ITS algorithm increased with tree height. This was expected because the larger trees are more clearly visible in ALS data. The best algorithm overall was AMS3D, with precision 0.5 and recall 0.41 for the tallest trees (>25 m). All algorithms tended to slightly underestimate the number of canopy trees and dramatically underestimate the number of understory trees.

In Sepilok, Dalponte2016+, AMS3D and Li2012 increased in precision and recall with tree height. AMS3D had the highest F1 score for the tallest trees (F1 = 0.71, 55-65 m). We note that the > 65 m class only contained 3 trees so we report accuracies for the penultimate height bin. Dalponte2016+ and AMS3D showed moderate accuracy (mean F1 = 0.3 and 0.49 respectively) for the medium sized trees (25-55 m), while Li2012 performed poorly in this range (mean F1 = 0.11) and was only accurate for trees over 55 m. The performance of Dalponte2016 was low for both understory and canopy trees, and moderate for medium sized trees (F1 = 0.42 for 25-35 m trees).

### 3.3 All ITS algorithms fail to segment understory trees

We tested the range of input parameters for each algorithm and we report the results for the parameter set with the highest accuracy. All ITS algorithms performed poorly for understory trees in both Sepilok (<25m, see Fig.5) and Wytham (<15m, see Fig.6). All of the algorithms had precision and recall scores below 0.1 for understory trees, with the exception of Dalponte2016 in Sepilok, which had a moderate precision (0.33) but very low recall (0.01) and F1 score (0.01). Importantly, the Wytham benchmark data set has a low pulse density, so understory trees have fewer ALS points, meaning that they are very challenging to accurately segment. This is representative of many ALS data sets. The Sepilok benchmark data set has a high pulse density and many understory trees with good coverage, which were nevertheless poorly segmented.

The reason for this poor performance in Dalponte2016, Dalponte2016+ and Li2012 is that they predicted too few trees in these low height classes. In Sepilok, the benchmark data contained 153 trees < 25 m, while Dalponte2016 predicts 17, Dalponte2016+ predicted 45 and Li2012 predicts 16 respectively. In Wytham, the benchmark data contained 11 trees < 15 m, while Dalponte2015 predicted 0, Dalponte2016+ predicts 1 and Li2012 predicts 0. AMS3D fails for the opposite reason: it predicted large numbers of very small trees in the low height classes, which do not match any of the trees in the benchmark data. Specifically, the most accurate predictions from AMS3D predicted 728 trees < 25 m in Sepilok and 132 trees <15 m in Wytham.

### 3.4 Sensitivity of segmentation accuracy to allometric parameters

The two best algorithms, AMS3D and Dalponte2016+ both have an input parameter related to the tree height to crown diameter allometry (H2CD and tau, respectively). This enables them to look for trees with larger crowns in the higher parts of the canopy. Unsurprisingly, the segmentation accuracy was highly sensitive to these parameters.

The accuracy of Dalponte2016+ was highest with tau = 80, where the search window size varies according to the the 80th percentile of the tree height to crown diameter allometry. The accuracy increased steadily from tau = 10 to tau = 80 and then dropped dramatically after this point (see figure 7). A similar pattern was observed in Wytham, with precision, recall and F1 score increasing with tau within from 10% to 70%. The accuracies then plateaued until 90% and finally decreased at 99%. This drop in accuracy for large tau was particularly noticeable in the medium size trees in both sites. Dalponte2016+ failed to detect <25m trees in Sepilok and <15m trees in Wytham regardless of tau, while it had decent precision, recall and F1 score when faced >55m and >25m with F1 score can reach 0.57(Sepilok) and 0.42(Wytham) respectively.

**Fig. 7.**
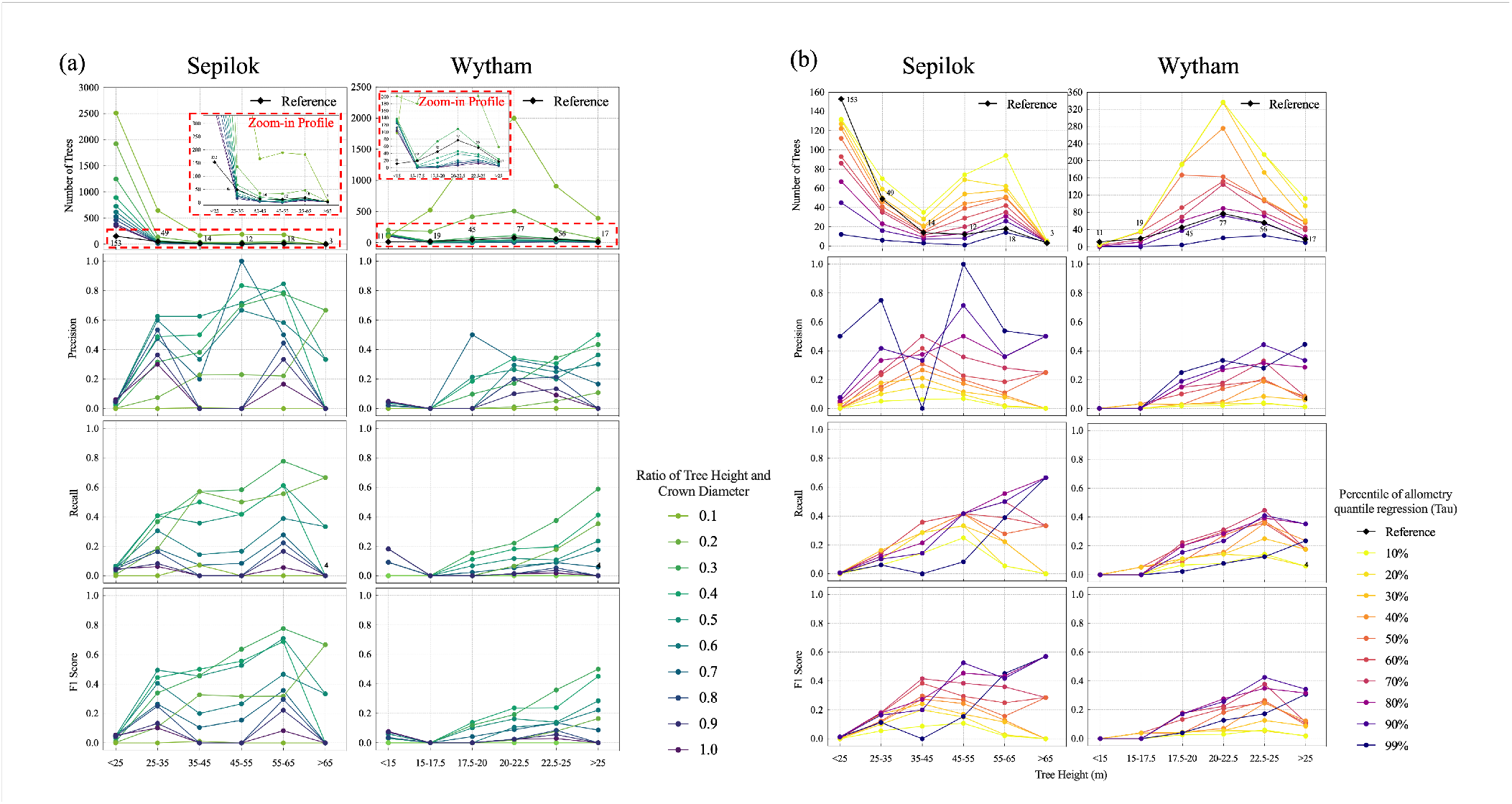
Allometric relation’s contributions to AMS3D and Dalponte2016+; The left Panel (a) shows the effects of tree height and crown diameter (H2CD) on AMS3D in Sepilok and Wytham plots; The right panel (b) displays the effects of tau on Dalponte2016+ in Sepilok and Wytham plots

The accuracy of AMS3D varied dramatically with the H2CD parameter, which controls the ratio between tree height and crown diameter. The accuracy was highest in with H2CD at 0.5 for Sepilok and 0.3 for Wytham. Variations in H2CD reduced this accuracy (see figure 7).

### 3.5 The number of trees predicted varies with allometric parameters

The number of trees predicted by each algorithm was highly sensitive to the allometric parameter (tau or H2CD). These algorithms can therefore be tuned to predict a realistic number and size distribution of trees, but this will reduce the overall accuracy of tree segmentation.

In Dalponte2016+, a higher tau led to fewer predictions throughout height ranges in both plots. Underestimation of the number of understory trees occurred for all allometry percentiles (closest estimate was 132 out of 146 <25m trees in Sepilok and 4 out of 9 <15m trees in Wytham). Dalponte2016+ overestimated the number of medium and large size trees when choosing <60% allometry percentiles in two sites.

In Sepilok, no clear trend was found as most allometry percentiles tended to underestimate the number of understory trees while overestimate medium and large trees. Among these allometry percentiles, 90% could be a better one if precision, recall and F1 score were also taken into consideration. A more complete visualization can be found in Figure I.1. of Supplementary I. In Wytham, Dalponte2016+ with 90% allometry percentile had predicted the number of trees >35m most accurately (182/193), though it still underestimated the <35m trees. A more complete visualization can be found in Figure I.2. of Supplementary I.

## 4 Discussion

### 4.1 Value of ALS benchmark data sets

ITS algorithms can accurately segment trees in conifer plantations but perform poorly for all but the canopy trees in forests with more complex structure. Tropical forests, which store huge amounts of carbon, therefore present an important challenge for ITS algorithms. It is critical that tree segmentation accuracy is assessed directly, rather than by comparing an ecological index such as biomass or the number and size distribution of trees, and benchmark data are needed to achieve this. Benchmark data sets are rare in broadleaf forests (although see (61)), especially in the tropics. Our Sepilok benchmark dataset is, to the best of our knowledge, the only available ITS benchmark data set in a tropical forest. The benchmark data sets used in this study (provided online) are particularly valuable as they include many understory trees. This was possible because we labelled the individual trees in the ALS data using TLS data, which contains detailed information on understory trees. We controlled for data quality issues by manually assigning a confidence score to each tree. This enables direct accuracy assessment for the understory trees, whereas most manually interpreted benchmark data sets focus only on the easily visible canopy trees. We hope other researchers use these benchmark data and follow our approach to produce additional benchmark sites, to help assess the accuracy of ITS algorithms across the tropics.

### 4.2 Tree segmentation algorithms are only accurate for canopy trees

We compared the performance the four most cited individual tree segmentation algorithms for ALS data in 1 ha of temperate and 1 ha of tropical forest. Our most striking result was that tree segmentation accuracy increased dramatically with tree height across both sites. This is likely because the top of the forest canopy is clearly visible to aerial surveys, while understory trees are often obscured. All algorithms, including 3D-point-cloud algorithms, failed to accurately segment understory trees. Note that we tested multiple combinations of input parameters for all algorithms (see Supplementary E and F), and the reported results from the parameter combination with highest accuracy. We note that the ALS pulse density was substantially lower in Wytham and therefore the understory trees contained fewer points, making segmentation more challenging. In Sepilok, many understory trees contained > 100 points and had a high confidence score.

### 4.3 Tree segmentation algorithms are more accurate when tuned to the local allometry

The most accurate ITS algorithm was AMS3D, closely followed by Dalponte2016+. The key similarity between these two algorithms is that they both contain a parameter which describes the expected relationship between tree height and crown size. This allometric information is widely available (53) and we suggest that users choose ITS algorithms which incorporate this information.

The value of including allometric information is clearly demonstrated by the fact that Dalponte2016+ dramatically outperformed Dalponte2016. These two algorithms both initially detect treetops before ‘growing’ the tree crowns until they reach certain stopping conditions. The only difference between them is that Dalponte2016+ detects treetops using a searching window whose width increases with canopy height. This enables it to look for trees with large crowns in tall areas of forest, and trees with small crowns in short areas of forest. Although allometric information can increase accuracy, it may also give false confidence if segmentation accuracy is not assessed directly. In Sepilok, we found that AMS3D could be tuned to predict roughly the correct number and size distribution of trees (with H2CD parameter = 0.9). However, the majority of these predicted trees did not correspond to real trees in the ground truth data set. The best performance was achieved with (H2CD = 0.5), and the understory trees were not well segmented by any combination of parameters. We therefore advise caution when assessing ITS algorithm accuracy, especially if the algorithm has been tuned to a local allometry.

### 4.4 What next for tree segmentation algorithms?

Understory trees are not visible in most remote sensing data, so can only be segmented using LiDAR data, which we show is currently inaccurate. This challenge may be addressed using extremely high pulse density LiDAR data collected using unoccupied Areial Vehicles (UAVs) (62). The processing power required to apply existing ALS segmentation algorithms to these massive data sets may be prohibitive, so we expect efficient algorithms to be adapted for this specific purpose.

Machine learning methods are proving helpful for distinguishing trees with studies ranging from object detection (63) to segmentation (64) and even species classification (65). One key challenge in this area is collecting sufficient reference data to train the machine learning models. We suggest that transfer learning could be helpful to lower the training costs (66, 67). Transfer learning has been successfully applied to 2D object detection (68, 69) and remote sensing-based landscape classification (70).

Recent studies have demonstrated the exciting potential of combining the structural information from ALS with spectral information to improve segmentation accuracy for canopy trees (71, 72). RGB or hyperspectral data can help distinguish tree crowns by their colour or texture, which are often species specific (73). This may therefore prove particularly useful in highly diverse tropical forests. We expect that combining techniques in this way will result in more reliable segmentation for the visible canopy trees.

## 5 Conclusions

This study compared the accuracy of four representative individual tree segmentation (ITS) algorithms against benchmark data sets in temperate and tropical forests. We found that all four ITS algorithms were able to segment canopy trees accurately but performed poorly for the understory trees. AMS3D was the most accurate ITS algorithm, closely followed by Dalponte2016+. Both of these algorithms benefited from using allometric information to improve their predictions. However, their accuracy was highly sensitive to these parameters. Crucially, we found that these parameters could be used to tune the algorithms to predict a realistic number and size distribution of trees, but that this actually reduced segmentation accuracy. This highlights the importance of robustly assessing segmentation results using labelled benchmark data sets, such as the openly available ones generated in this study.

## Data and code availability

Benchmark data for Sepilok Forest and Wytham Woods: https://zenodo.org/deposit/7181101

Code demo for labelling ALS with TLS: https://github.com/Elephant-C/ALS-labelling

Code demo for tree crown polygon-based assessment: https://github.com/Elephant-C/tree-crown-based-assessment

## Author contributions

TJ, YC and JB conceived the idea and designed the study. YC conducted all the analysis and prepared the code demo used in the study. LS and NK developed the AMS3D software. KC, AB, TJ, PW and MD collected the TLS data in both Wytham and Sepilok. PW and KC processed the TLS data in Sepilok and Wytham respectively. TJ and DC organized the ALS data collection in Sepilok. DC organized the ALS data collection in Wytham. TJ and YC led the writing of the manuscript, in close collaboration with JB. All authors contributed critically to the drafts and gave final approval for publication.

## Acknowledgements

YC was funded by China Scholarship Council. TJ and DC were funded by NE/S010750/1. The Sepilok ALS data collection was funded by NE/S010750/1. We thank Ground Data Solutions R & D Sdn Bhd for collecting the ALS data in Sepilok. We thank the Natural Environment Research Council Airborne Research Facility for collecting the Wytham ALS data. The Wytham TLS fieldwork was funded through the Metrology for Earth Observation and Climate project (MetEOC-2), grant number ENV55 within the European Metrology Research Programme (EMRP). The EMRP is jointly funded by the EMRP participating countries within EURAMET and the European Union. Funds for purchase of the UCL RIEGL VZ-400 instrument was provided by the UK NERC National Centre for Earth Observation (NCEO) and UCL Geography. We thank Niall Origo for the help with the Wytham fieldwork. K.C was funded by the European Union (ERC-2021-STG Grant agreement No. 101039795). Views and opinions expressed are however those of the author(s) only and do not necessarily reflect those of the European Union or the European Research Council Executive Agency. Neither the European Union nor the granting authority can be held responsible for them. YL was fund by Key Project of Shanghai Science and technology Innovation Action (No. 20dz1201202).

