## Supplementary for "Tree segmentation in airborne laser scanning data is only accurate for canopy trees": appendix.pdf

### A Reviews for State-of-the-art airborne LiDAR ITS algorithms and inter-comparison works

We reviewed state-of-the-art airborne LiDAR individual tree segmentation algorithms in terms of history citation, algorithm structure, forest type, experiment setting as well as main results. The details can be found at: ITS algorithm overview. Besides, we also did literature review about previously ITS inter-comparison work, and the corresponding details can be found at: ITS algorithm intercomparison overview.

### B Confidence score system and visualizations for ALS reference individual trees with confidence score

We introduced a confidence score system to data quality of ALS individual tree point clouds. The system was defined by manually checking the ALS individual tree point clouds with corresponding TLS point clouds. In general, there are three main categories (good/medium/bad) to describe quality of individual tree reference data, while it could be furtuher divided into 7 different levels in terms of data richness, point distribution and individual tree crown feature. In our study, we included good (Confidence score=7 and 6) and medium (Confidence Score=5) in the inter-comparison for both Sepilok and Wytham case as we were confident about their quality.

**Table B.1.** Definition of confidence score system

| Class | Confidence Score | Definition |
|---|---|---|
| Good | 7 | Enough ALS point clouds to fit tree crown polygon, and the individual tree feature representation is accurate |
| | 6 | Enough ALS point clouds to fit tree crown polygon, and the bias for individual tree feature representation is small |
| Medium | 5 | Enough ALS point cloud to fit tree crown polygons, but the bias for individual tree feature representation is larger |
| Bad/Died Tree | 4 | Enough ALS point clouds to fit tree crown polygons, but the number of points are not enough to represent a tree |
| | 3 | ALS data is available, but too few point clouds (less 3 points) to form tree crown polygons |
| | 2 | No ALS data available |
| | 1 | Tree has fallen down |

DRAFT

Figure B.1 below shows the tree counts with regards to tree height and confidence score for Sepilok Forest and Wytham Woods. The ALS individual tree reference visualizations are in the following section C and D

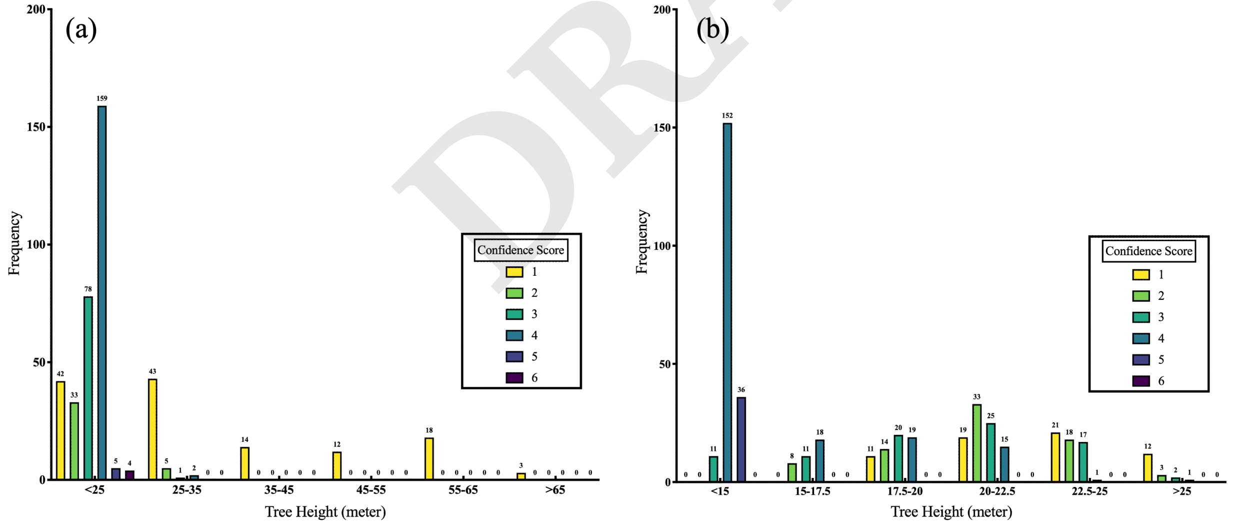


**Fig. B.1.** ALS reference tree frequencies with regards to tree height and confidence score; X axis refers to different tree height and Y axis means the number of trees (frequency); Different bars colorized with confidence score; Left Panel (a) shows details for Sepilok ALS reference; Right Panel (b) displays Wytham Woods ALS reference.

### C Visualization for Sepilok Forest ALS individual tree point clouds

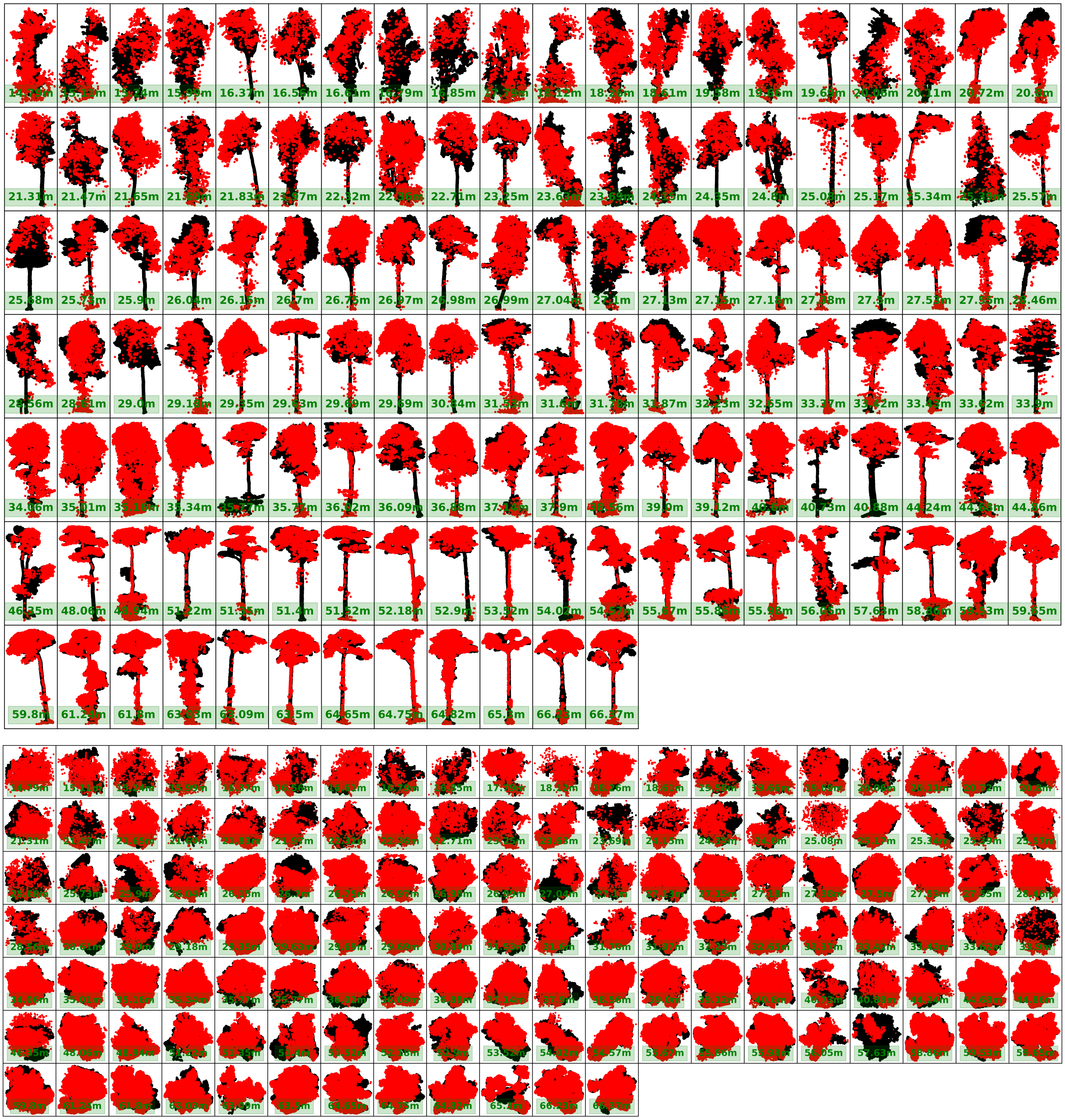

**Fig. C.1.** Visualization for individual trees from Sepilok forest case with confidence score = 7; Figures in upper panel display side view of individual tree and figures in the bottom panel are corresponding top-down view

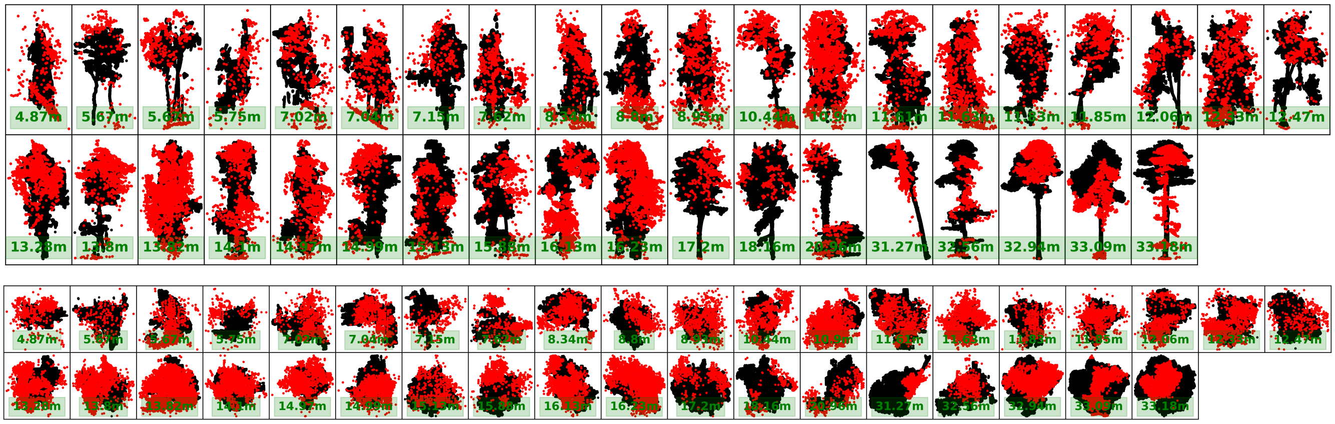

**Fig. C.2.** Visualization for individual trees from Sepilok forest case with confidence score = 6; Figures in upper panel display side view of individual tree and figures in the bottom panel are corresponding top-down view

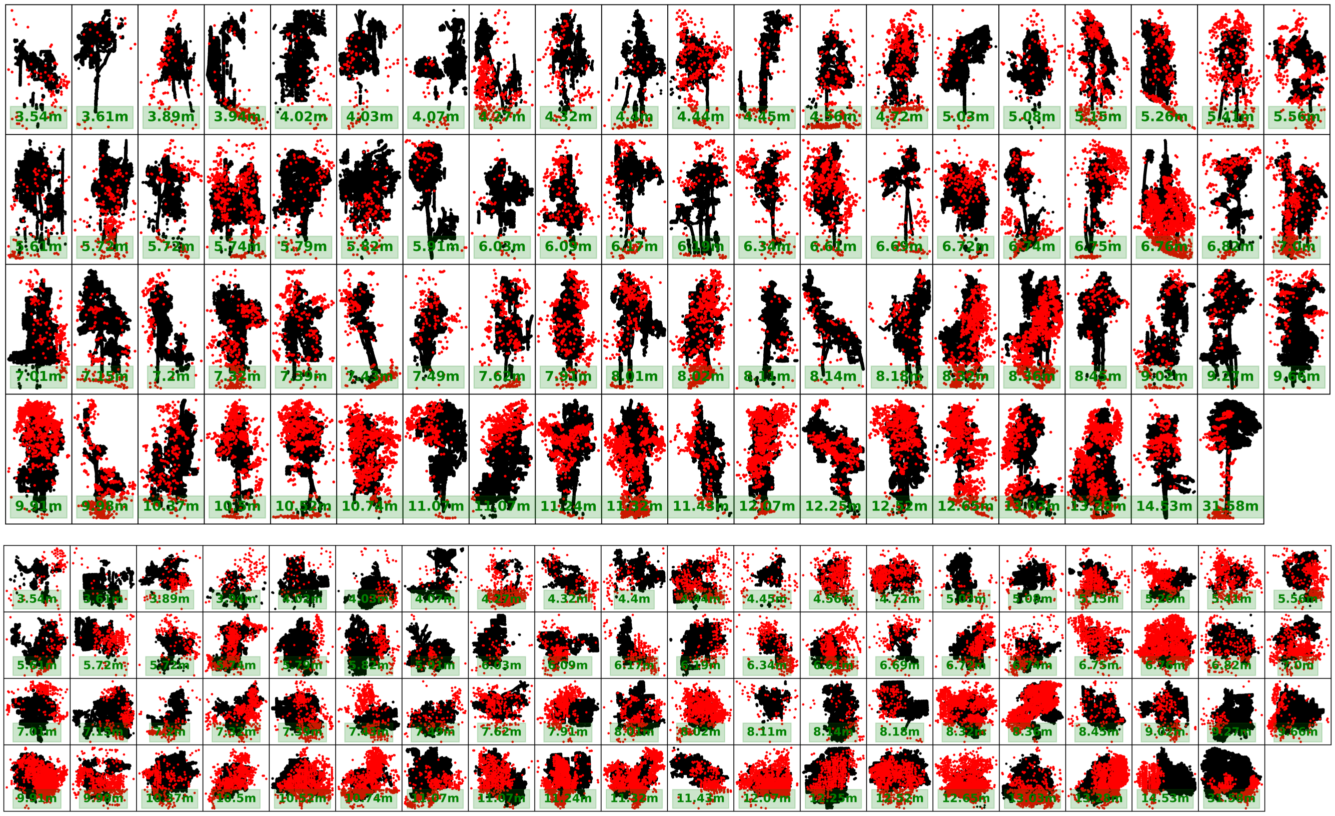

**Fig. C.3.** Visualization for individual trees from Sepilok forest case with confidence score = 5; Figures in upper panel display side view of individual tree and figures in the bottom panel are corresponding top-down view

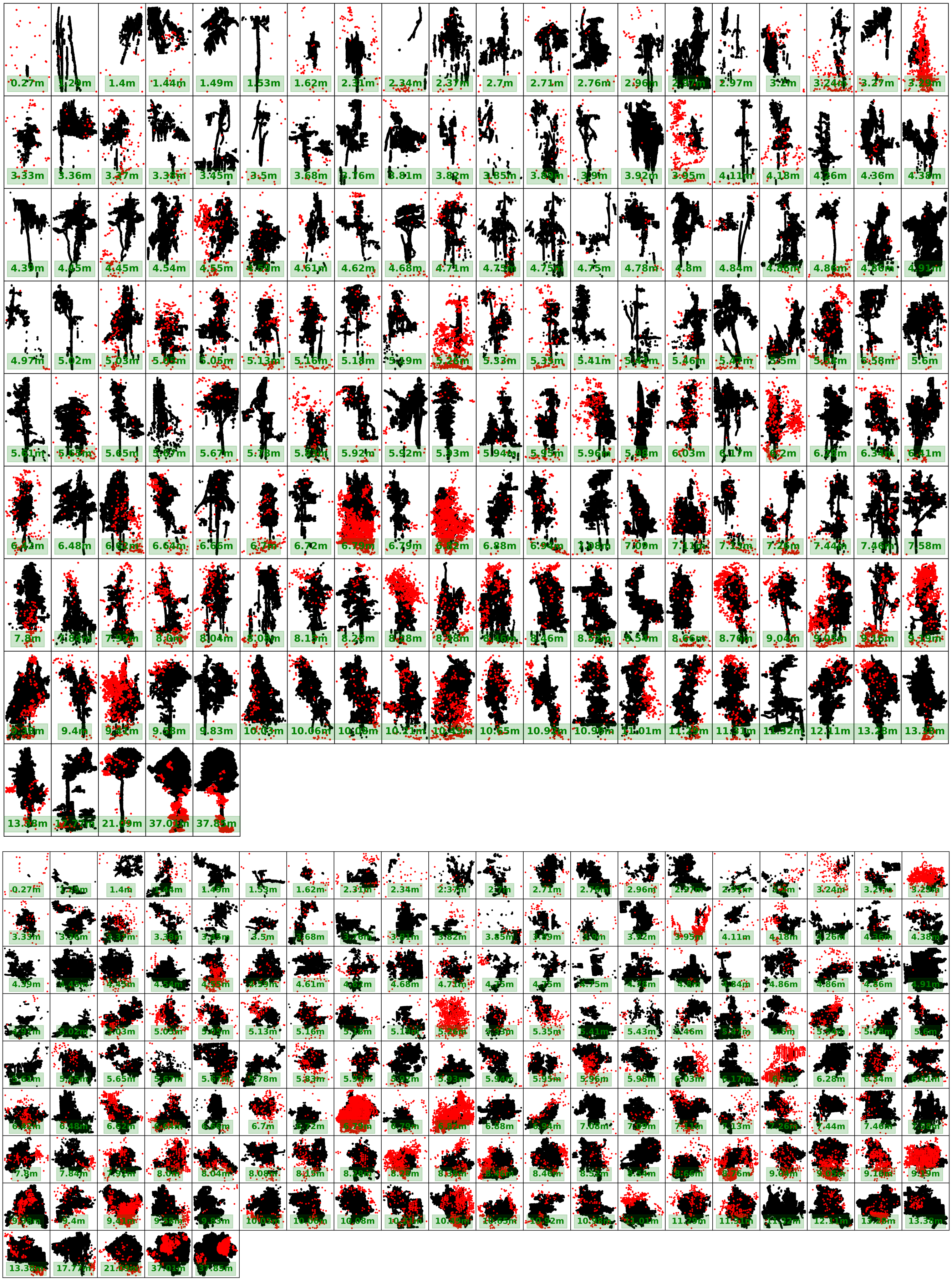

**Fig. C.4.** Visualization for individual trees from Sepilok forest case with confidence score = 4; Figures in upper panel display side view of individual tree and figures in the bottom panel are corresponding top-down view

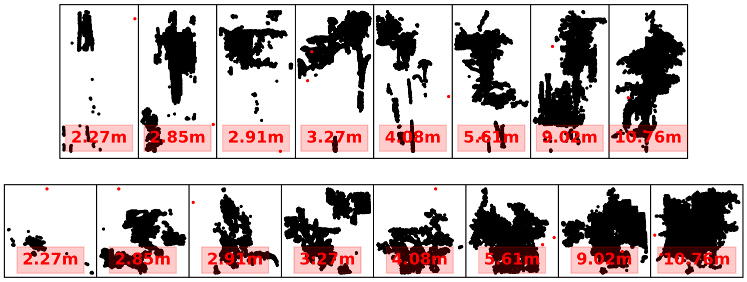


**Fig. C.5.** Visualization for individual trees from Sepilok forest case with confidence score = 3; Figures in upper panel display side view of individual tree and figures in the bottom panel are corresponding top-down view

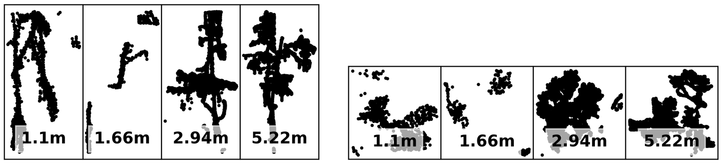


**Fig. C.6.** Visualization for individual trees from Sepilok forest case with confidence score = 2; Figures in left panel display side view of individual tree and figures in the right panel are corresponding top-down view

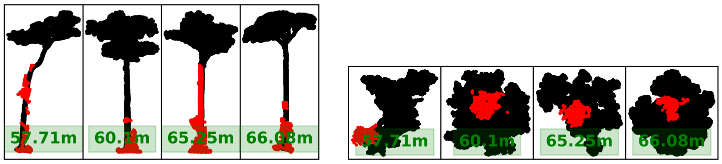

**Fig. C.7.** Visualization for individual trees from Sepilok forest case with confidence score = 1; Figures in left panel display side view of individual tree and figures in the right panel are corresponding top-down view

### D Visualization for Wytham Woods ALS individual tree point clouds

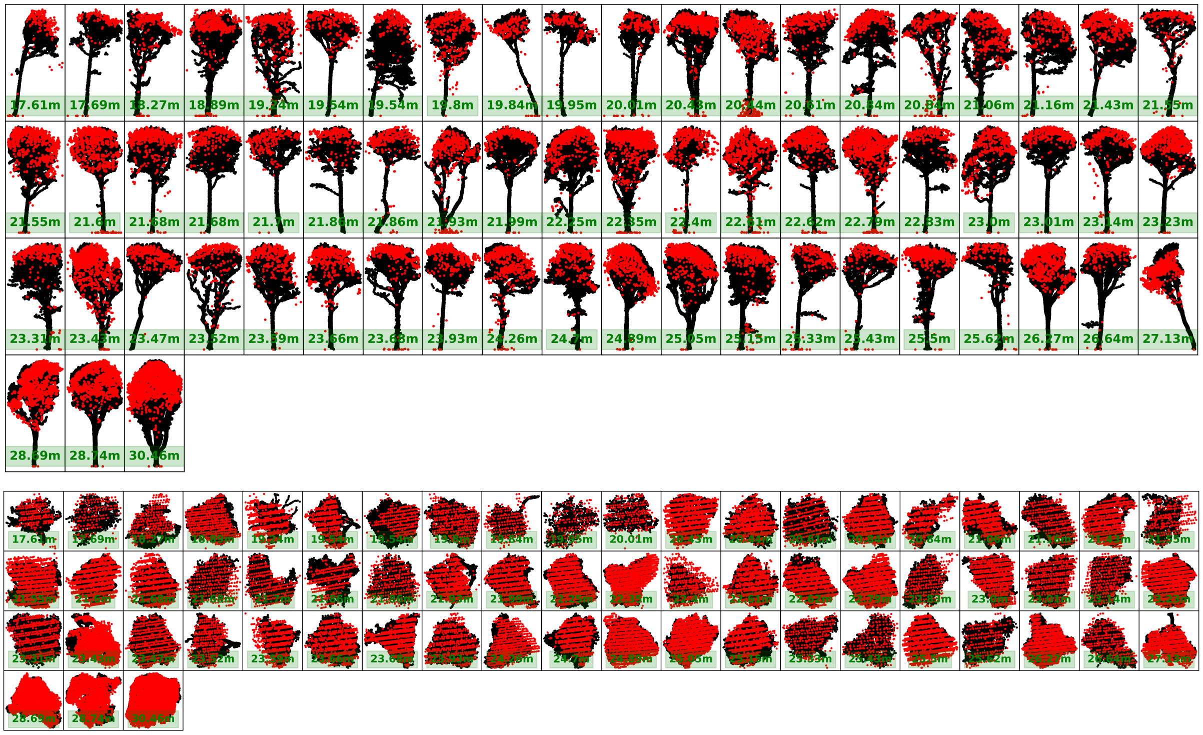

**Fig. D.1.** Visualization for individual trees from Wytham wood case with confidence score = 7; Figures in upper panel display side view of individual tree and figures in the bottom panel are corresponding top-down view

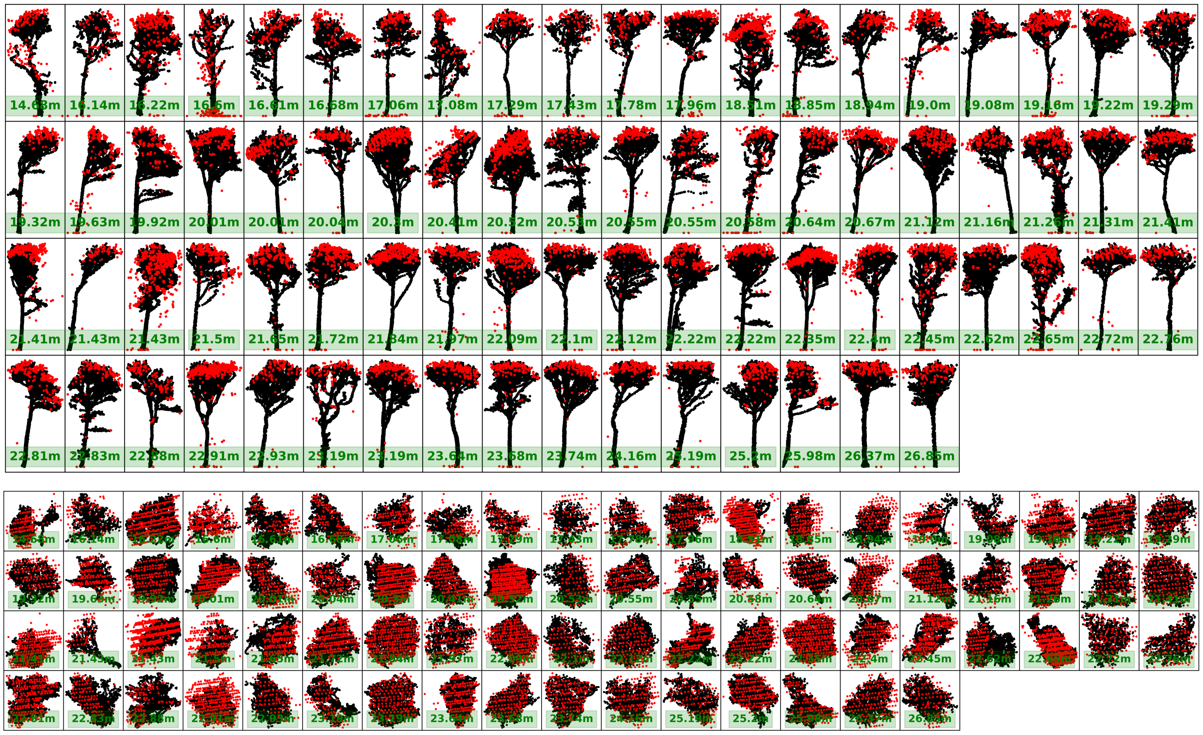

**Fig. D.2.** Visualization for individual trees from Wytham wood case with confidence score = 6; Figures in upper panel display side view of individual tree and figures in the bottom panel are corresponding top-down view

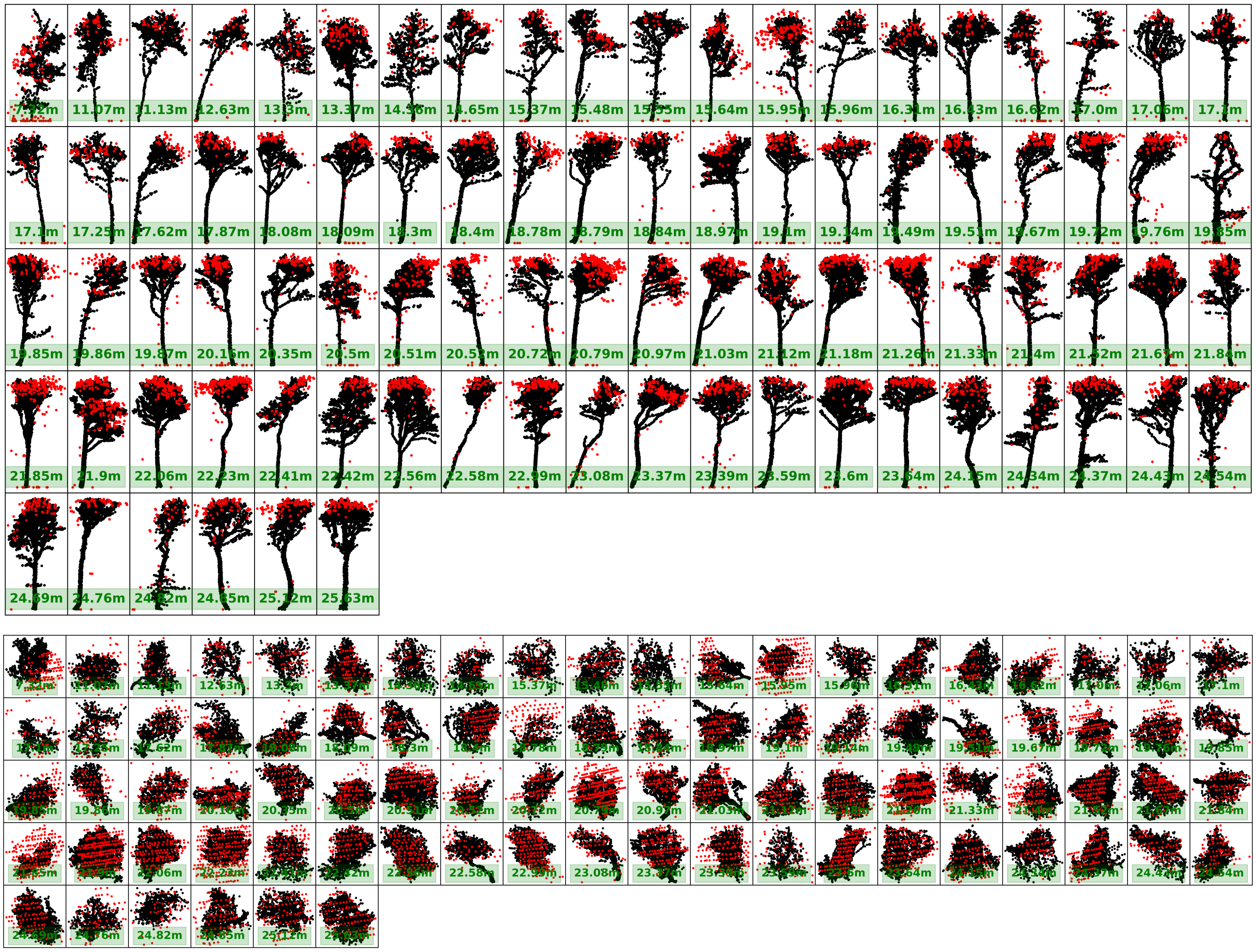

**Fig. D.3.** Visualization for individual trees from Wytham wood case with confidence score = 5; Figures in upper panel display side view of individual tree and figures in the bottom panel are corresponding top-down view

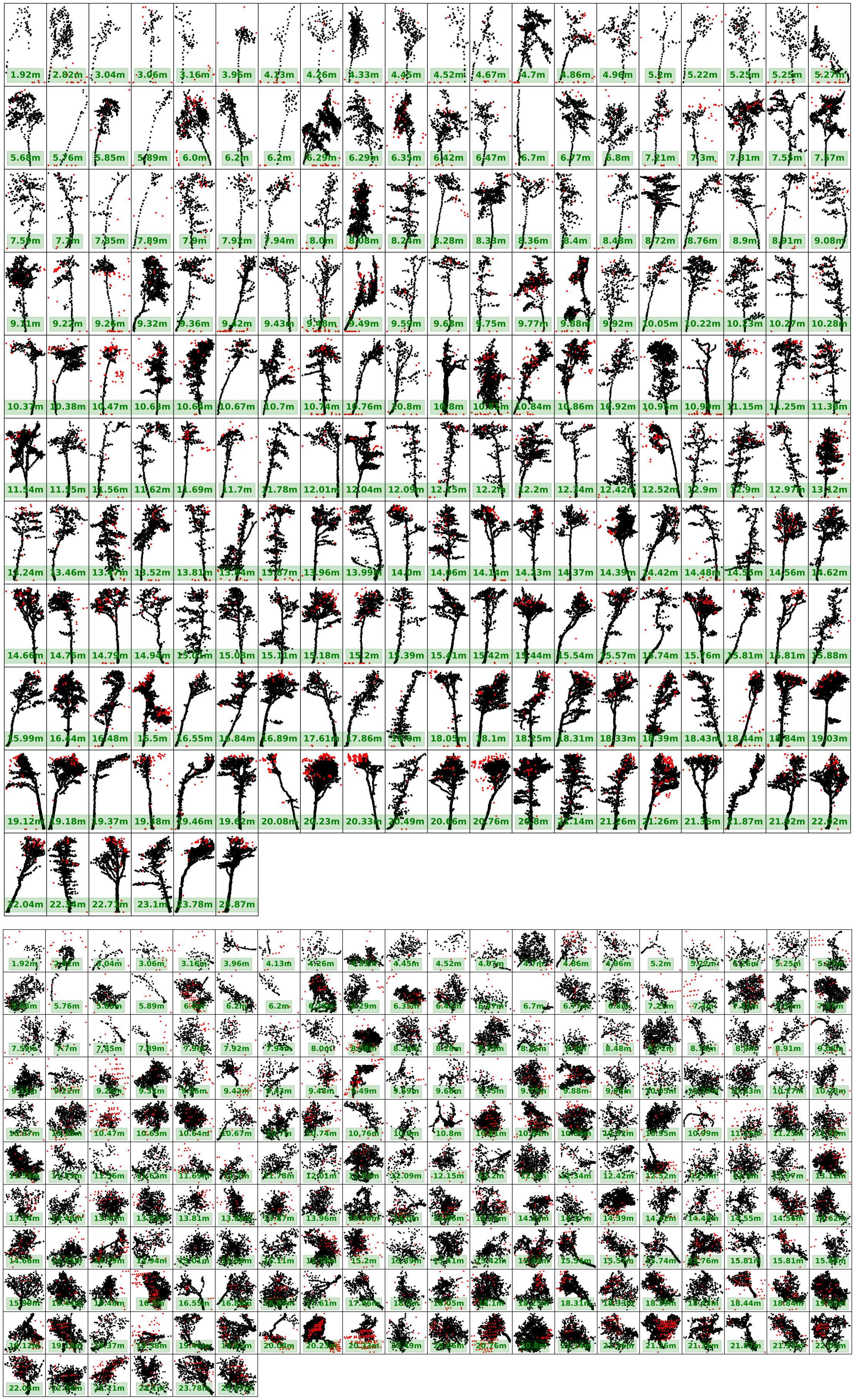

**Fig. D.4.** Visualization for individual trees from Wytham wood case with confidence score = 4; Figures in upper panel display side view of individual tree and figures in the bottom panel are corresponding top-down view

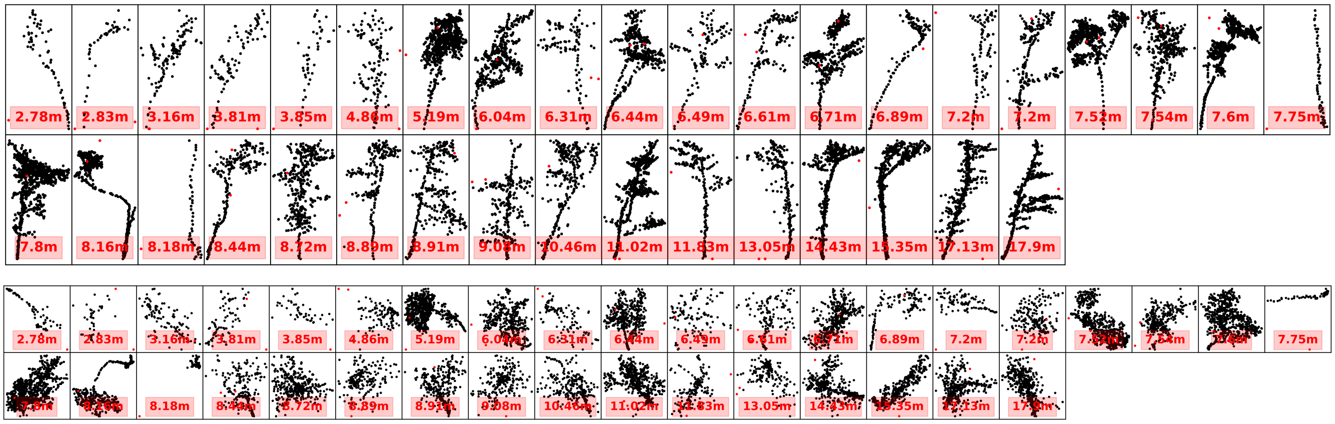


**Fig. D.5.** Visualization for individual trees from Wytham wood case with confidence score = 3; Figures in upper panel display side view of individual tree and figures in the bottom panel are corresponding top-down view

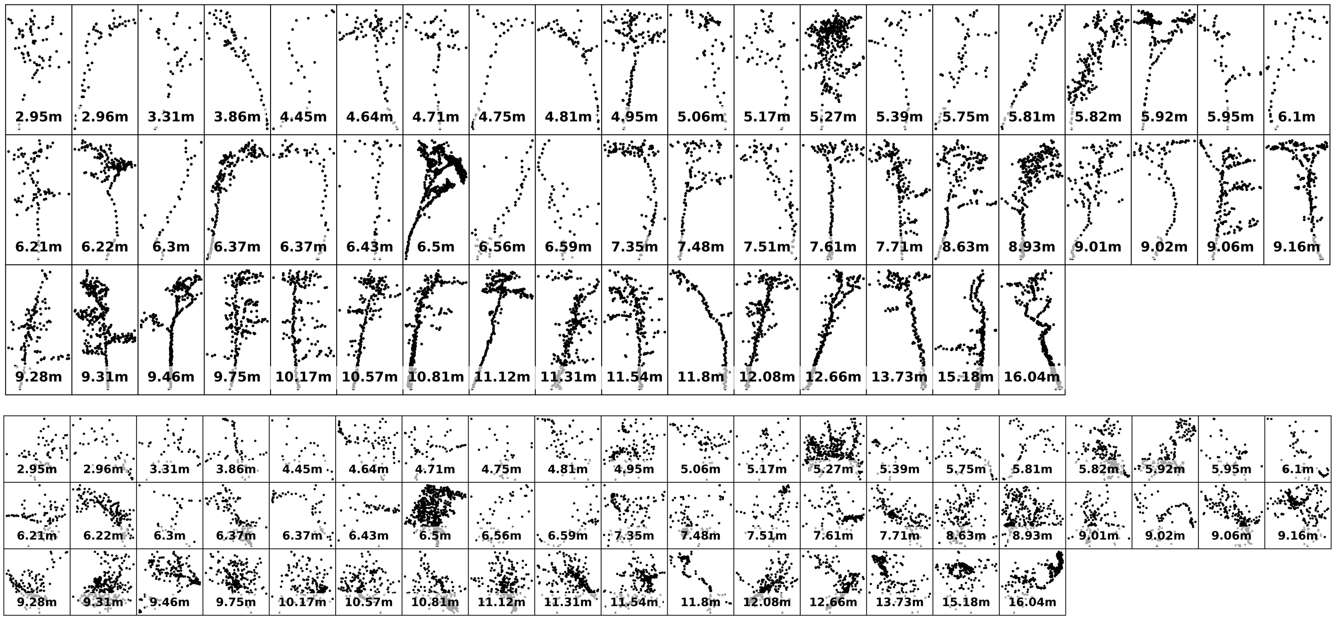


**Fig. D.6.** Visualization for individual trees from Wytham wood case with confidence score = 2; Figures in upper panel display side view of individual tree and figures in the bottom panel are corresponding top-down view

### E Matrix plots to show Precision, Recall and F1 of 4 ITS algorithms in Sepilok Forest with different Confidence Score Combinations

We followed the tree crown polygon-based assessment framework and implemented grid searching for the four ITS algorithm with corresponding paramter space introduced in main texts, and matrix plots for precision, recall and f1 score are shown as follows:

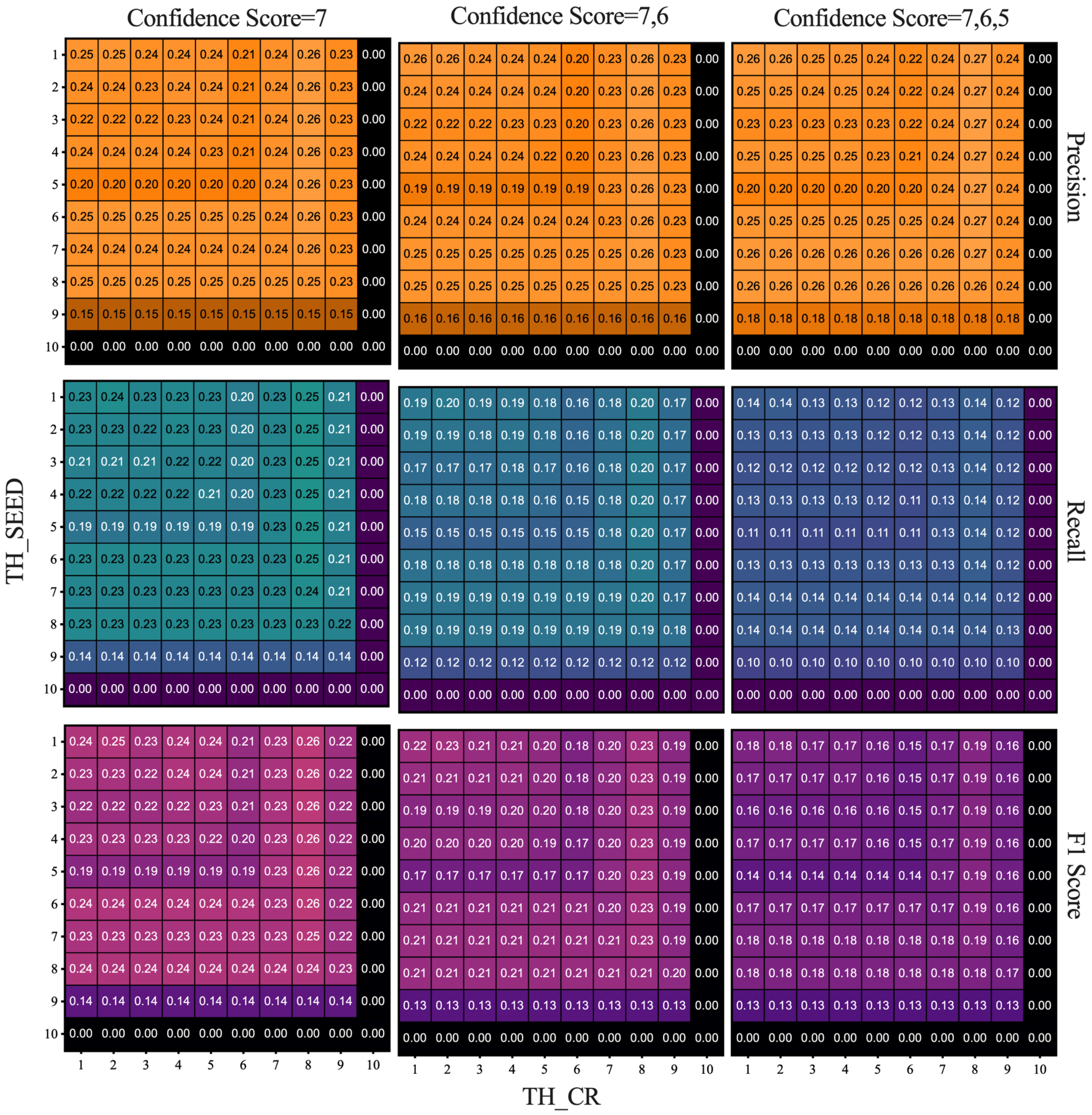


**Fig. E.1.** Precision, Recall and F1 Score matrix plots for Dalponte2016 in Sepilok Forest

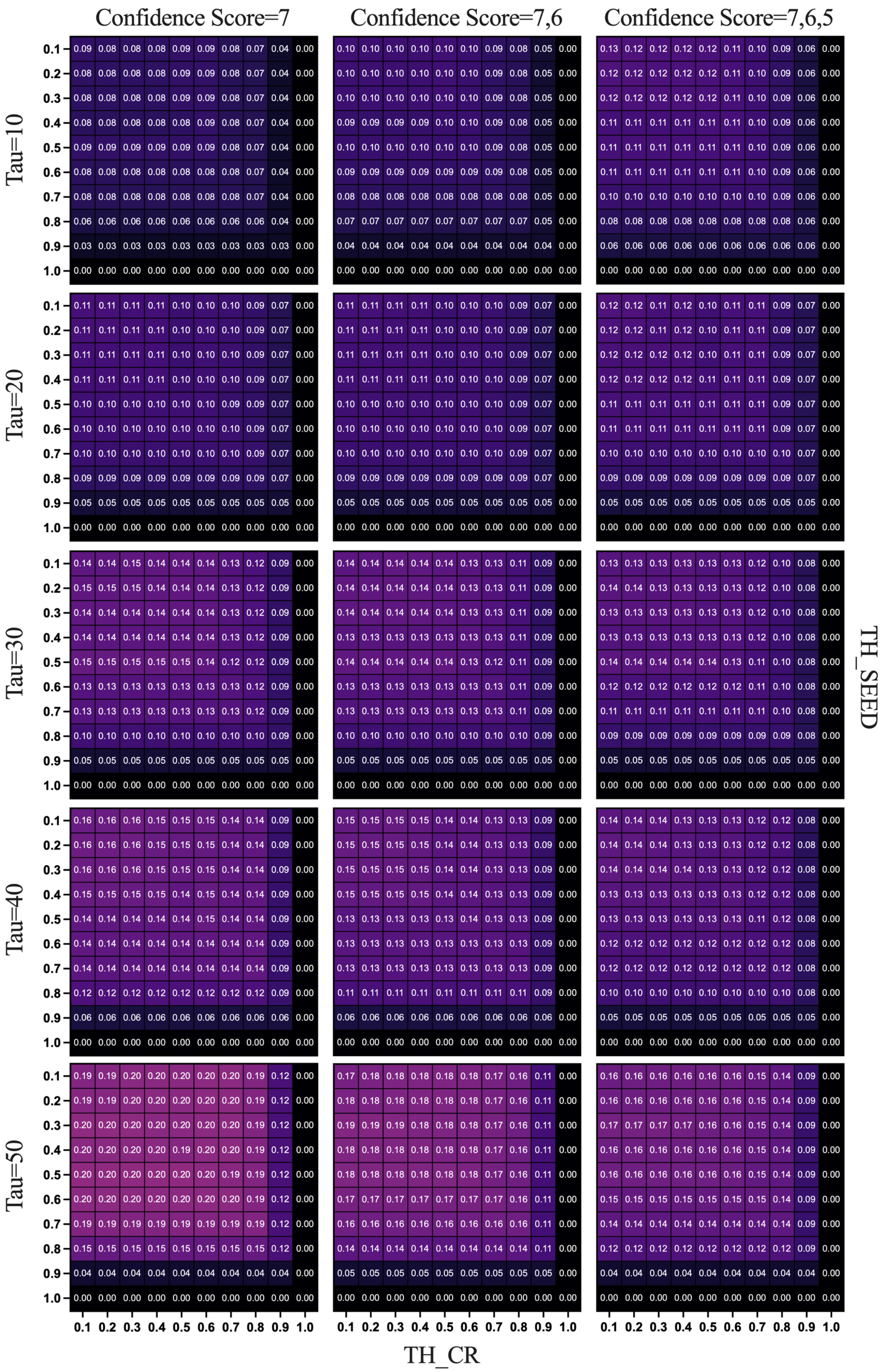


**Fig. E.2.** Precision, Recall and F1 Score with regards to Tau ranging from 10 to 50 percent matrix plots for Dalponte2016+ in Sepilok Forest

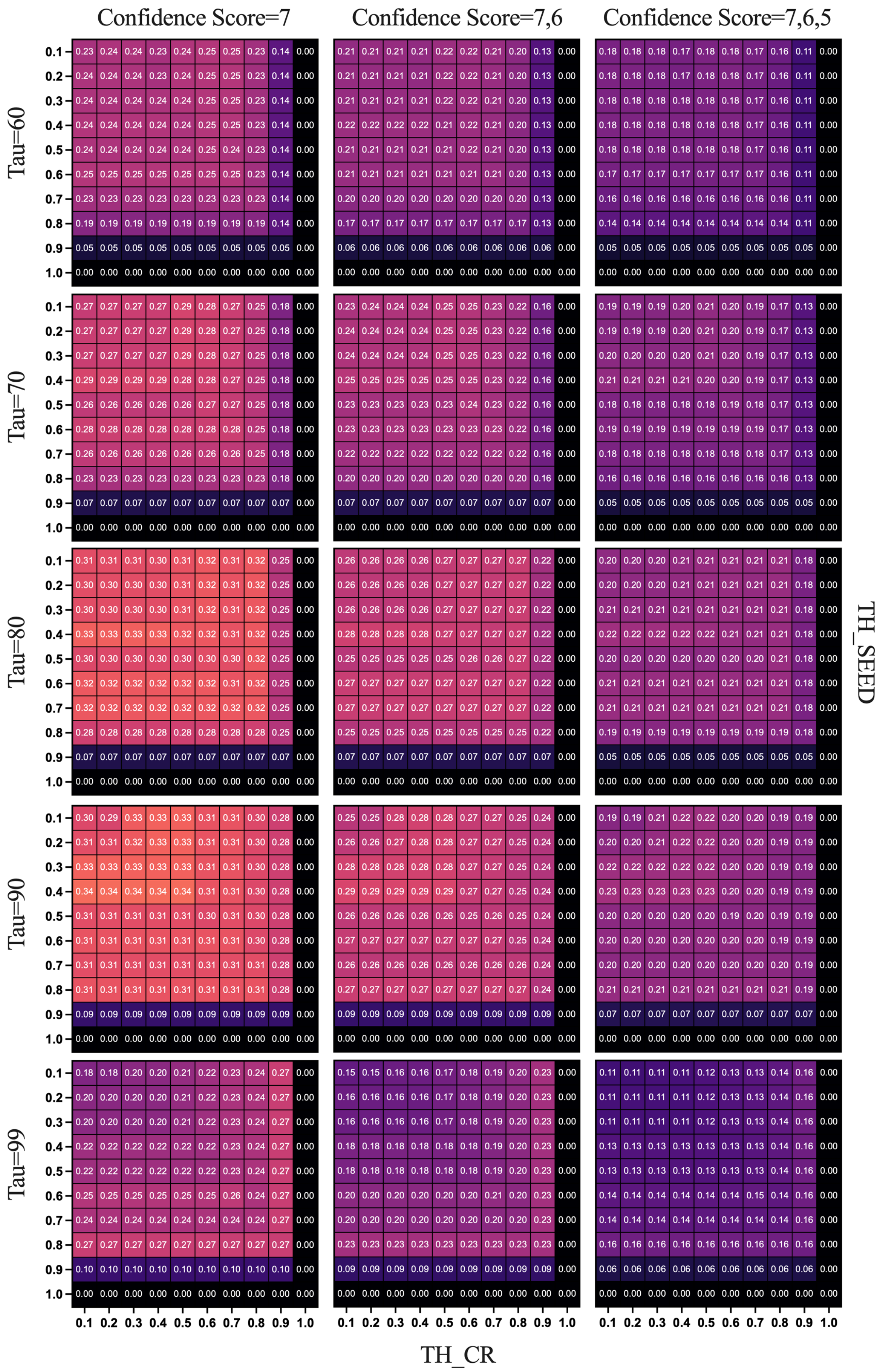


**Fig. E.3.** Precision, Recall and F1 Score with regards to Tau ranging from 60 to 90 percent matrix plots for Dalponte2016+ in Sepilok Forest

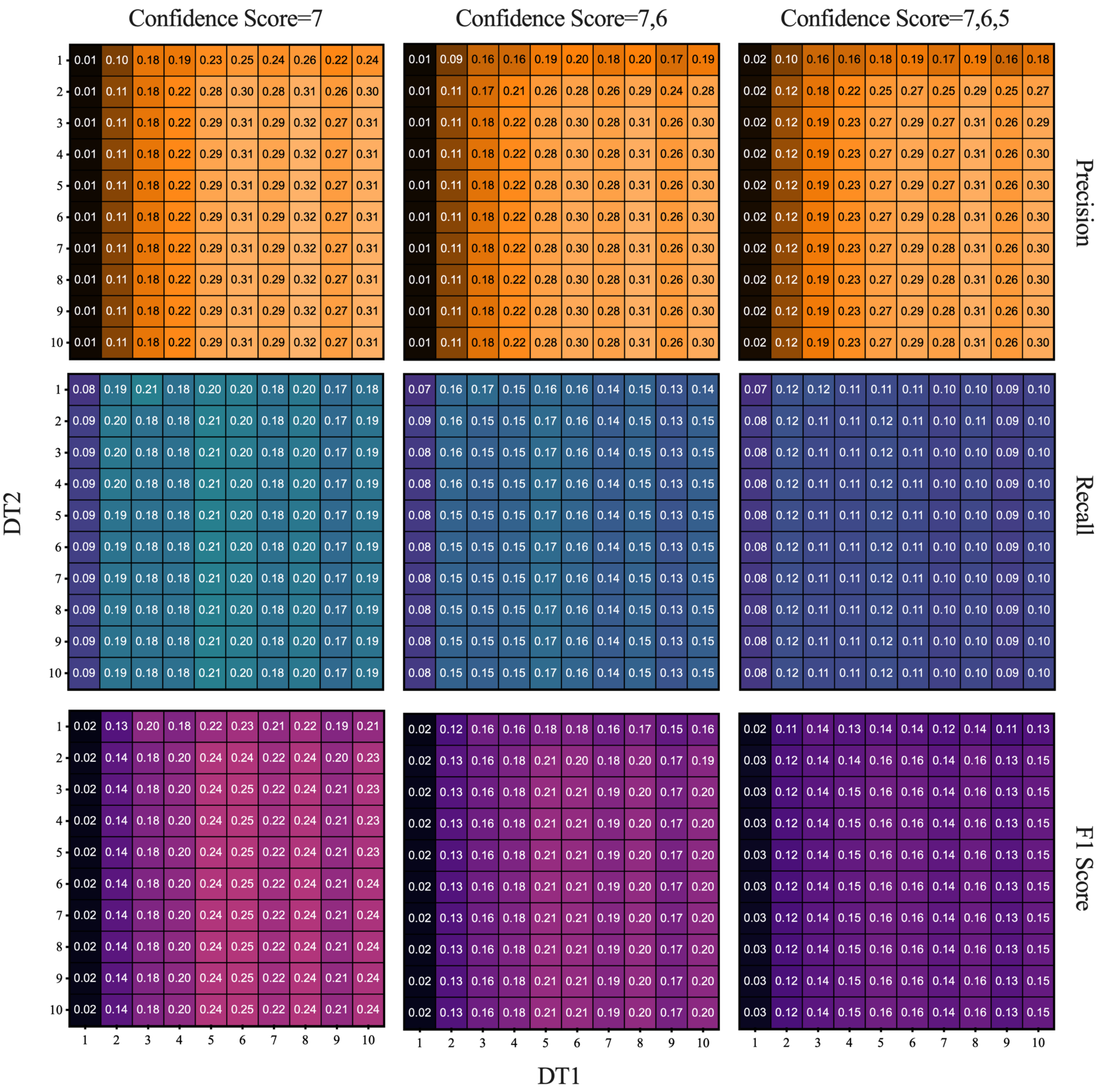


**Fig. E.4.** Precision, Recall and F1 Score matrix plot for Li2012 in Sepilok Forest

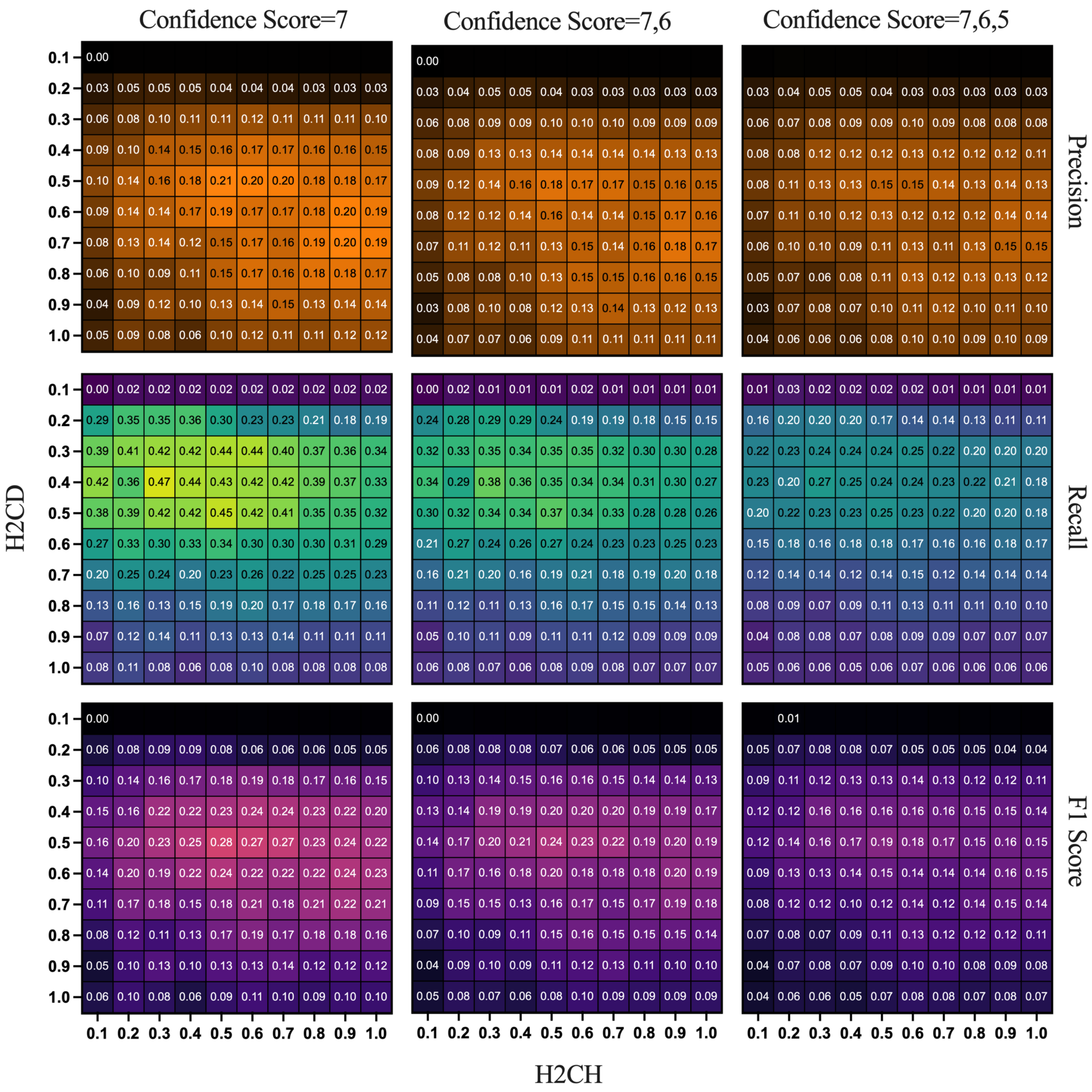


**Fig. E.5.** Precision, Recall and F1 Score matrix plot for AMS3D in Sepilok Forest

### F Matrix plots to show Precision, Recall and F1 of 4 ITS algorithms in Wytham Woods with different Confidence Score Combinations

Precicion, Recall and F1 Score matrix plots for the 4 ITS algorithms display as follows.

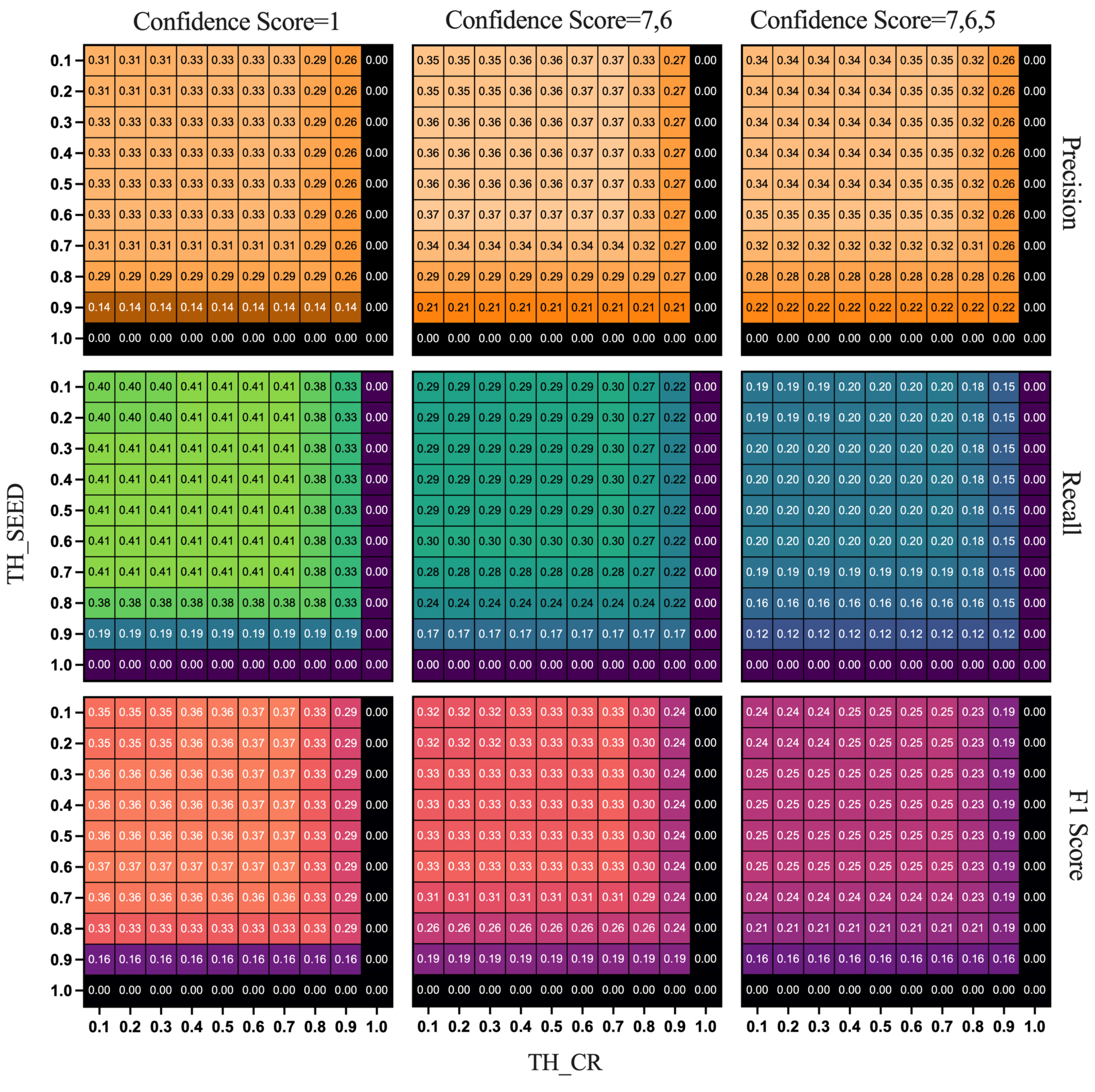


**Fig. F.1.** Precision, Recall and F1 Score matrix plot for Dalponte2016 in Wytham Woods

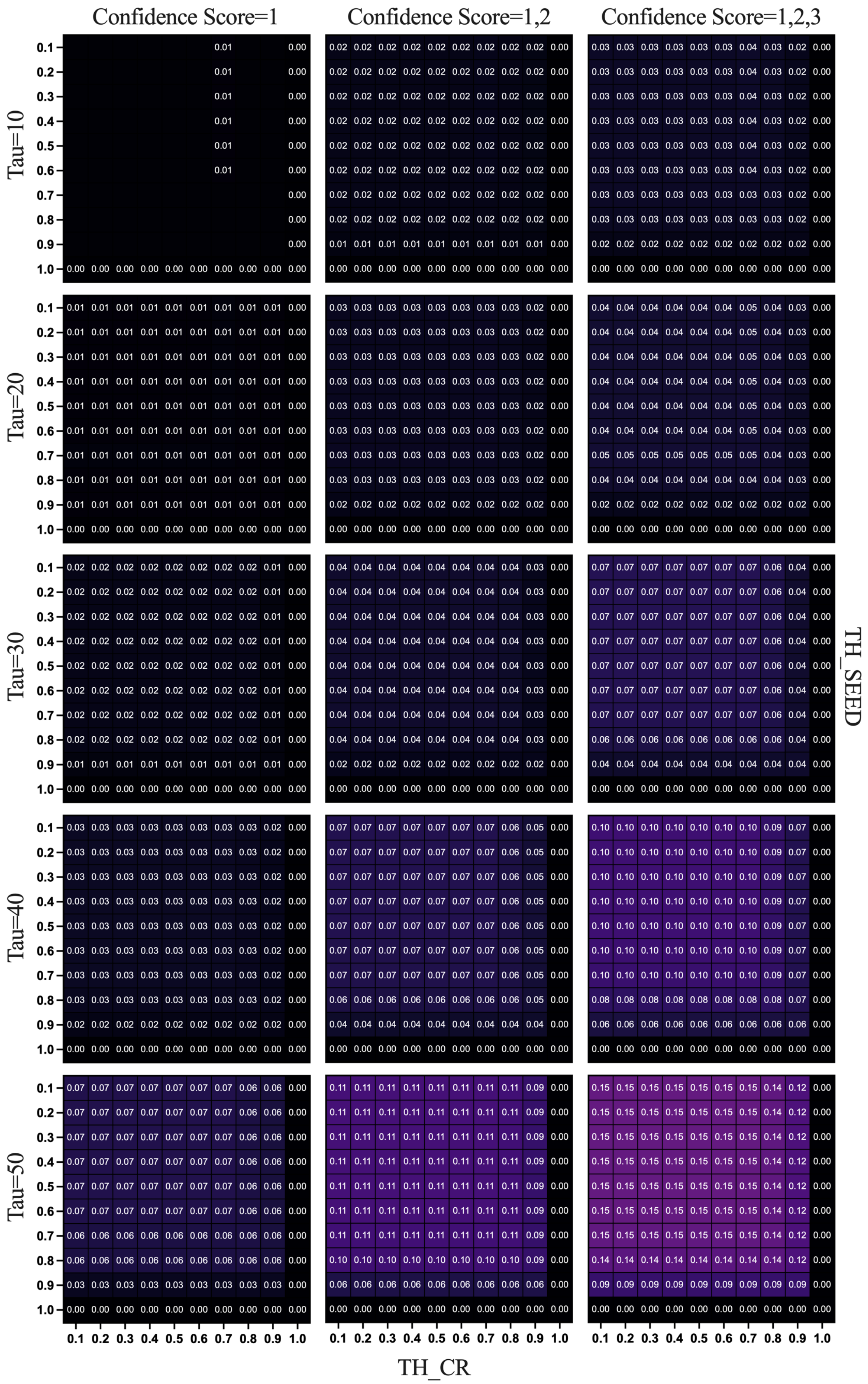


**Fig. F.2.** Precision, Recall and F1 Score with regards tau ranging from 10 to 50 percent matrix plot for Dalponte2016+ in Wytham Woods

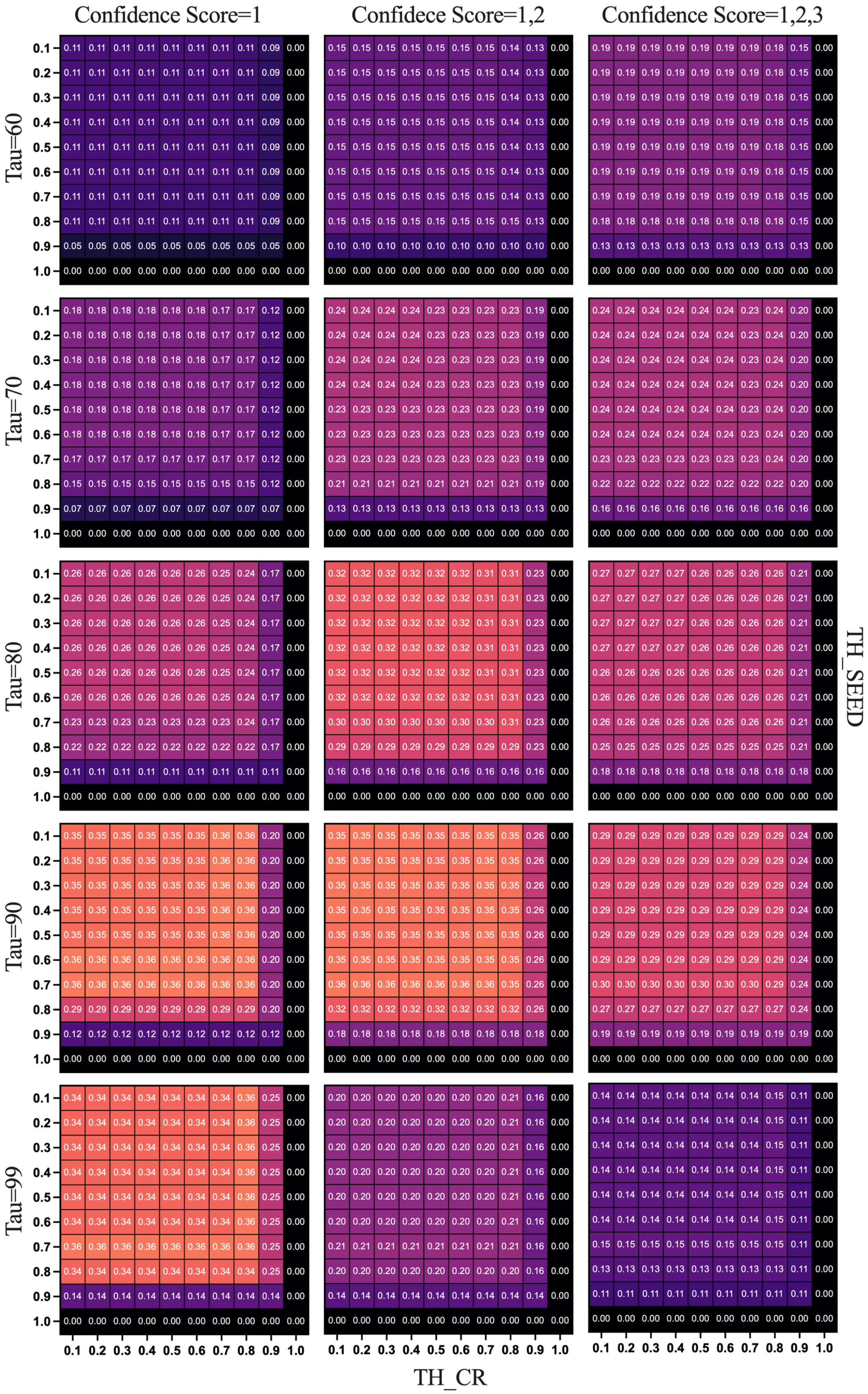


**Fig. F.3.** Precision, Recall and F1 Score matrix plot for Dalponte2016 in Wytham Woods

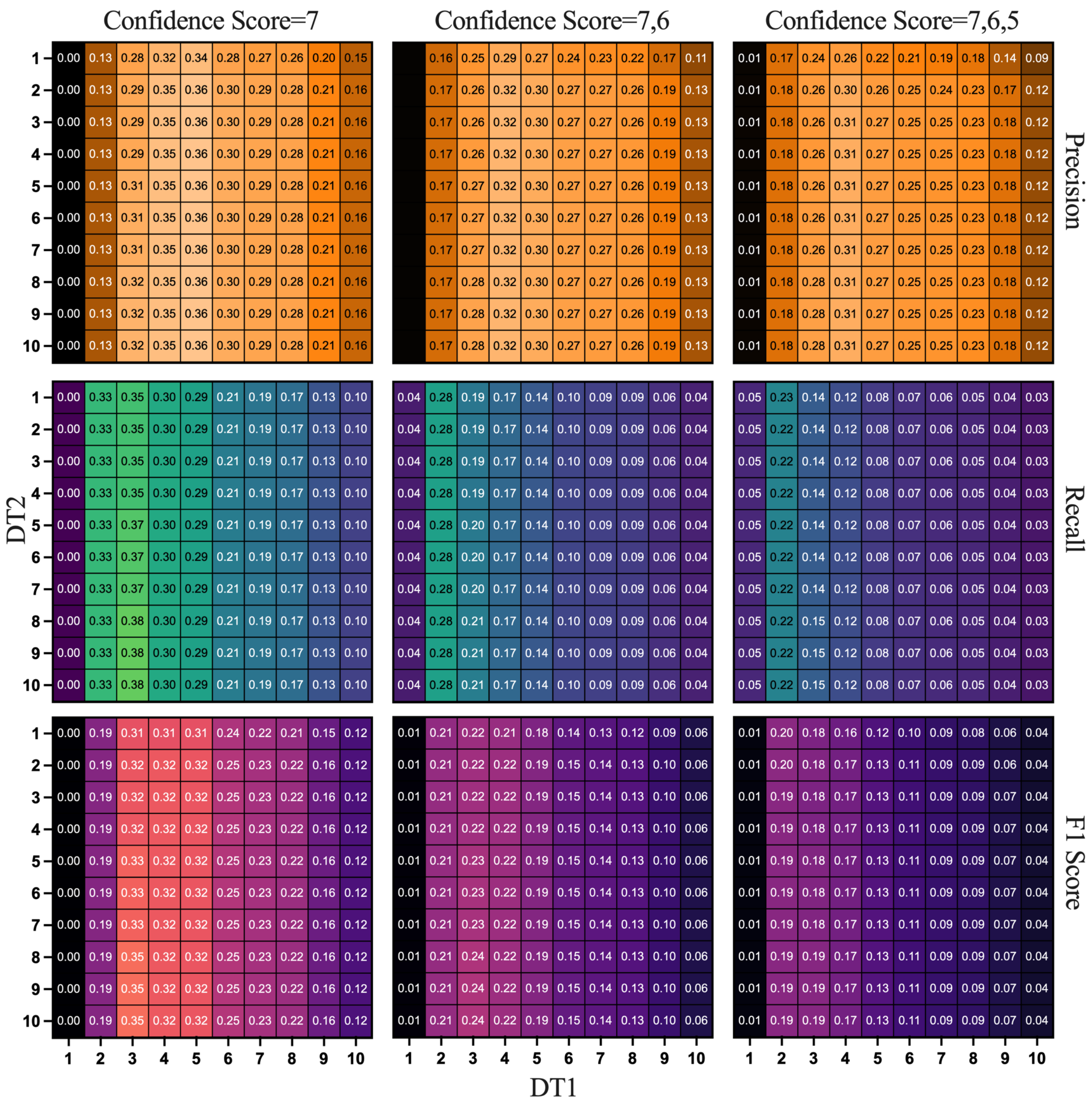


**Fig. F.4.** Precision, Recall and F1 Score matrix plot for Dalponte2016 in Wytham Woods

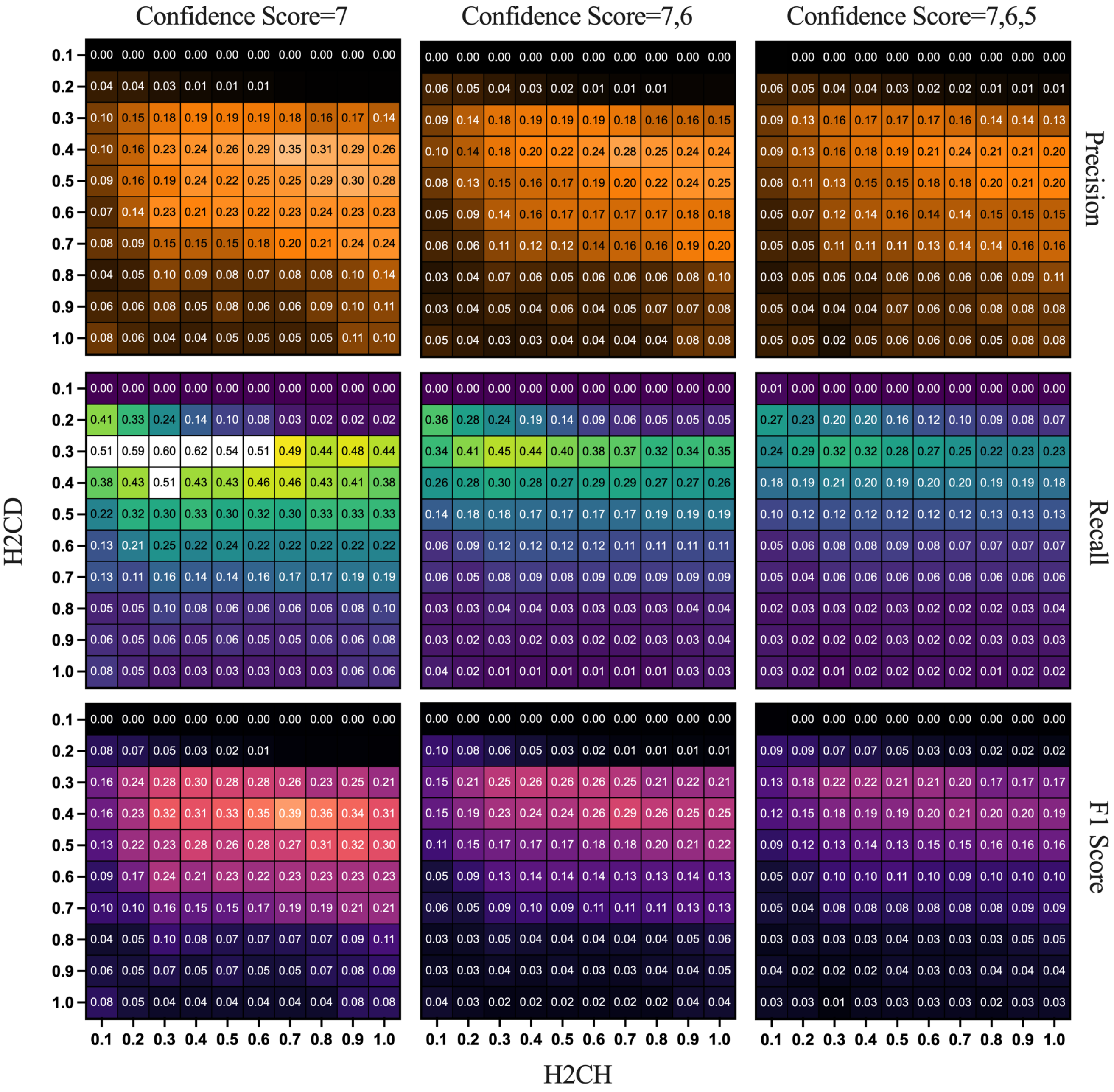


**Fig. F.5.** Precision, Recall and F1 Score matrix plot for Dalponte2016 in Wytham Woods

### G Precision, Recall and F1 Score with regards to height relations and multiple Confidence Score Combinations in Sepilok Forest

Precision, Recall and F1 changes regarding tree heights for 4 involved ITS algorithms in Sepilok Forest plot show as follows:

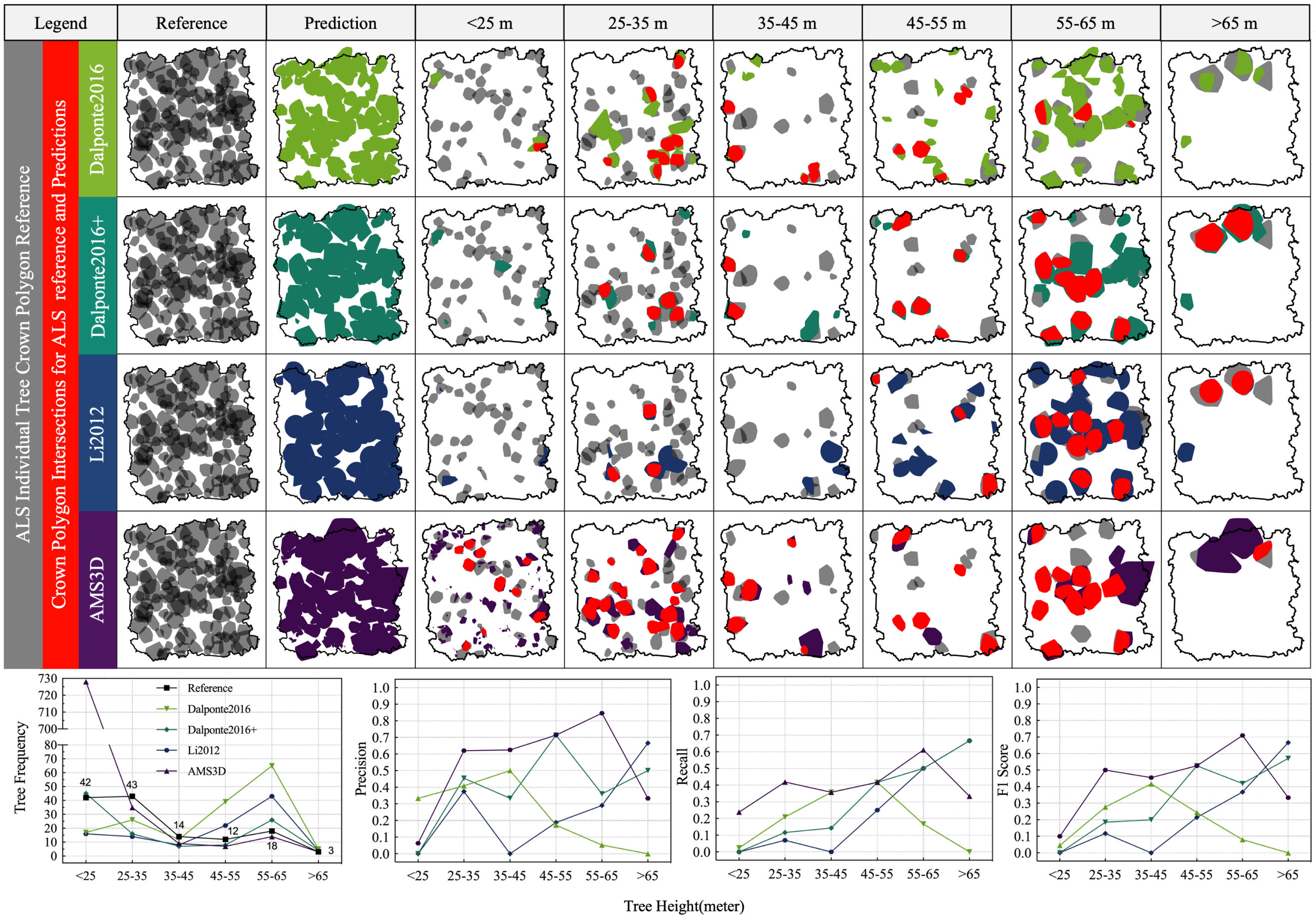


**Fig. G.1.** Precision, Recall and F1 Score changes with regards to tree height and confidence score=7 in Sepilok Forest

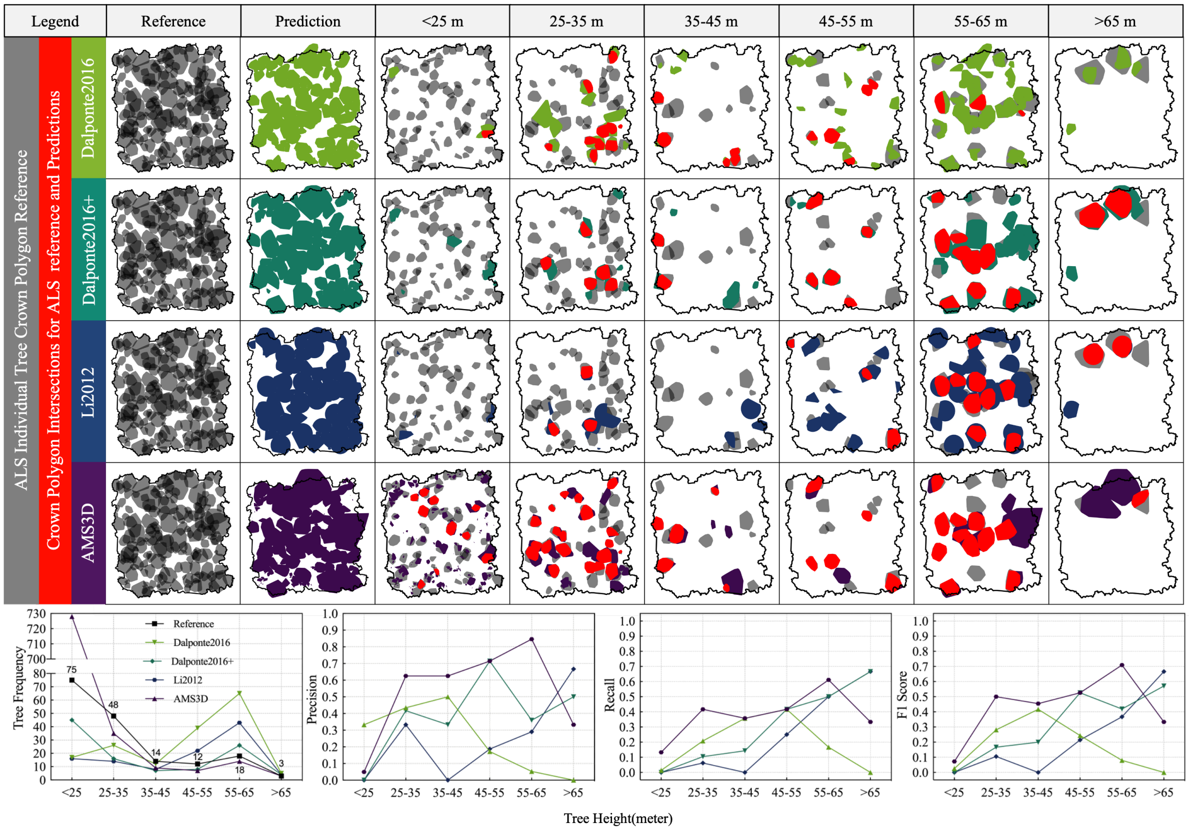


**Fig. G.2.** Precision, Recall and F1 Score changes with regards to tree height and confidence score=7,6 in Sepilok Forest

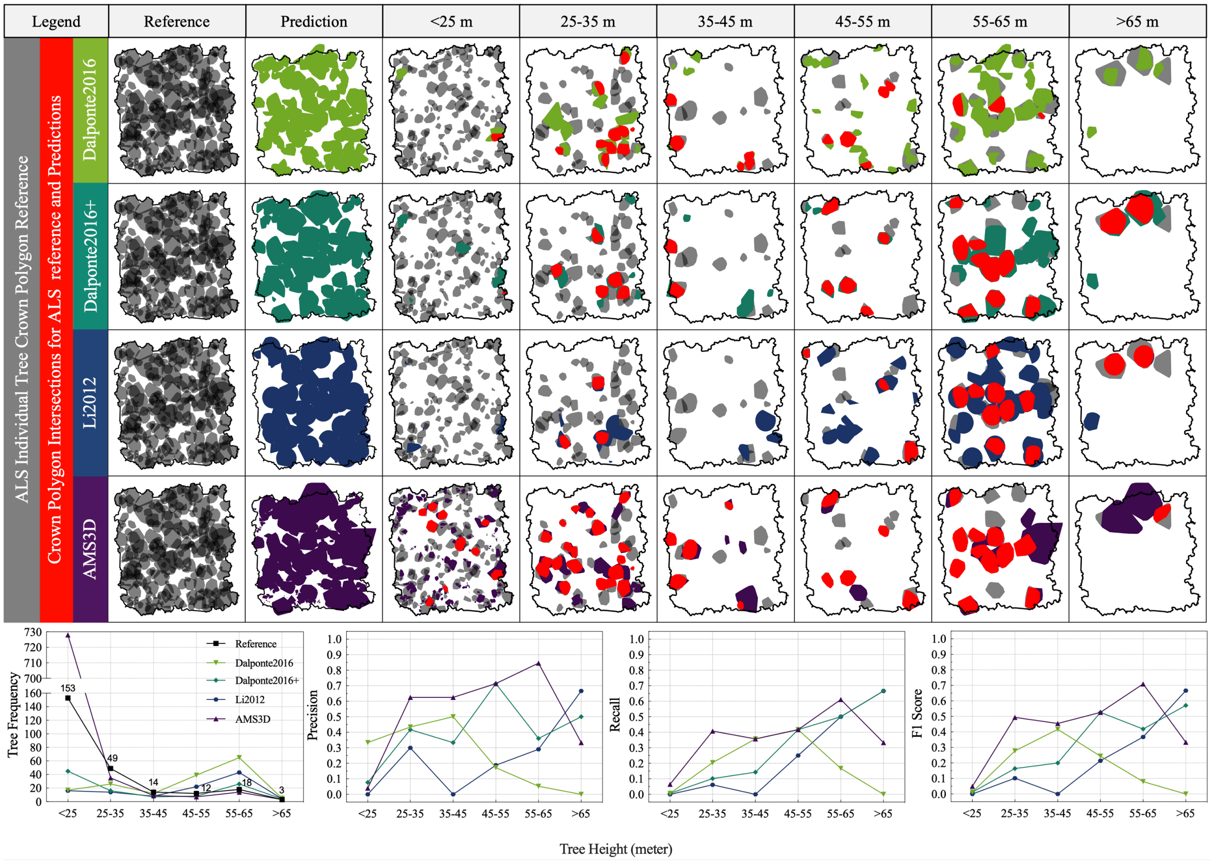


**Fig. G.3.** Precision, Recall and F1 Score changes with regards to tree height and confidence score=7,6,5 in Sepilok Forest

### H Precision, Recall and F1 Score with regards to height relations and multiple Confidence Score Combinations in Wytham Woods

Precision, Recall and F1 changes regarding tree heights for 4 involved ITS algorithms in Wyhtam Woods plot show as follows:

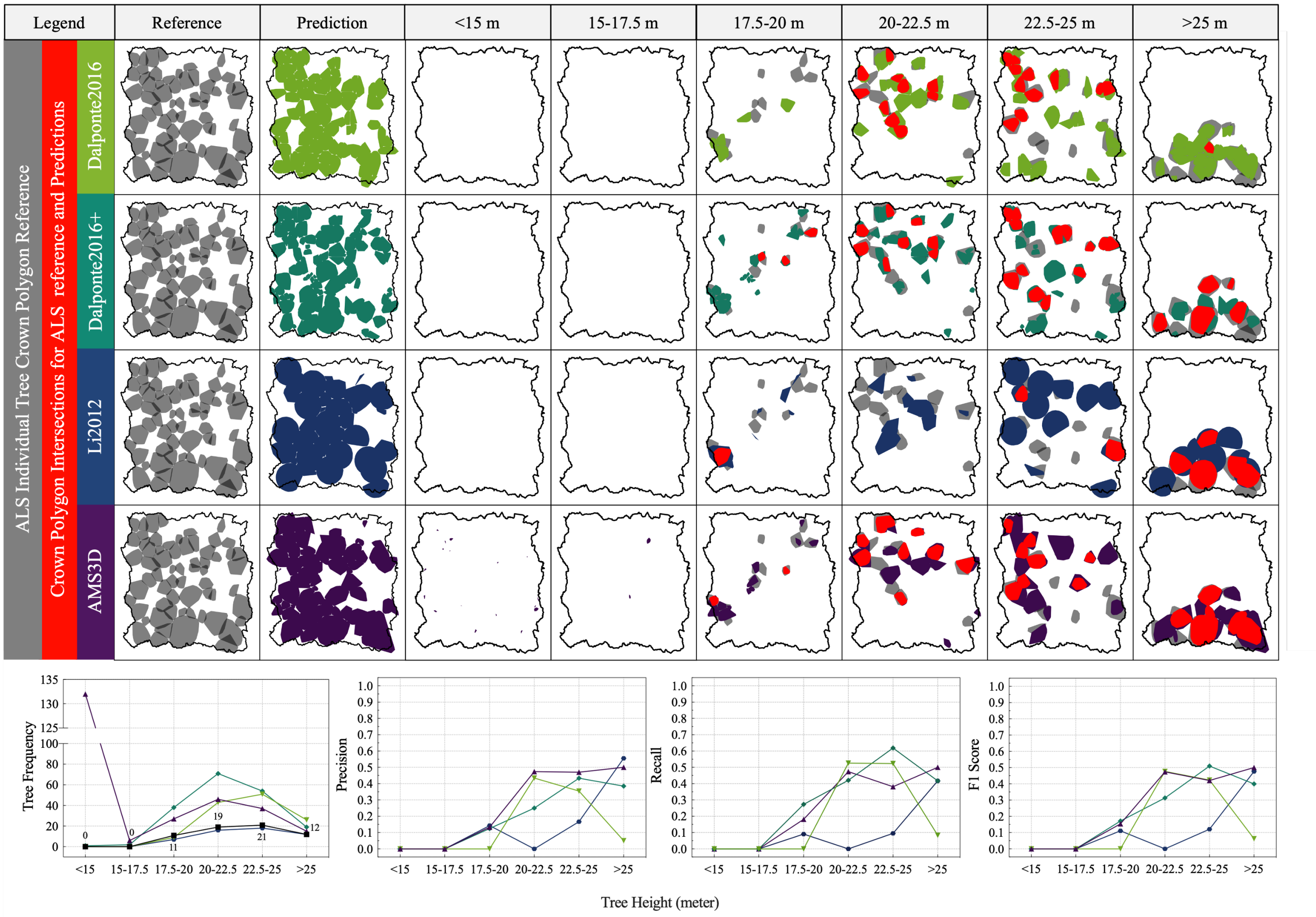


**Fig. H.1.** Precision, Recall and F1 Score changes with regards to tree height and confidence score=7 in Wytham Woods

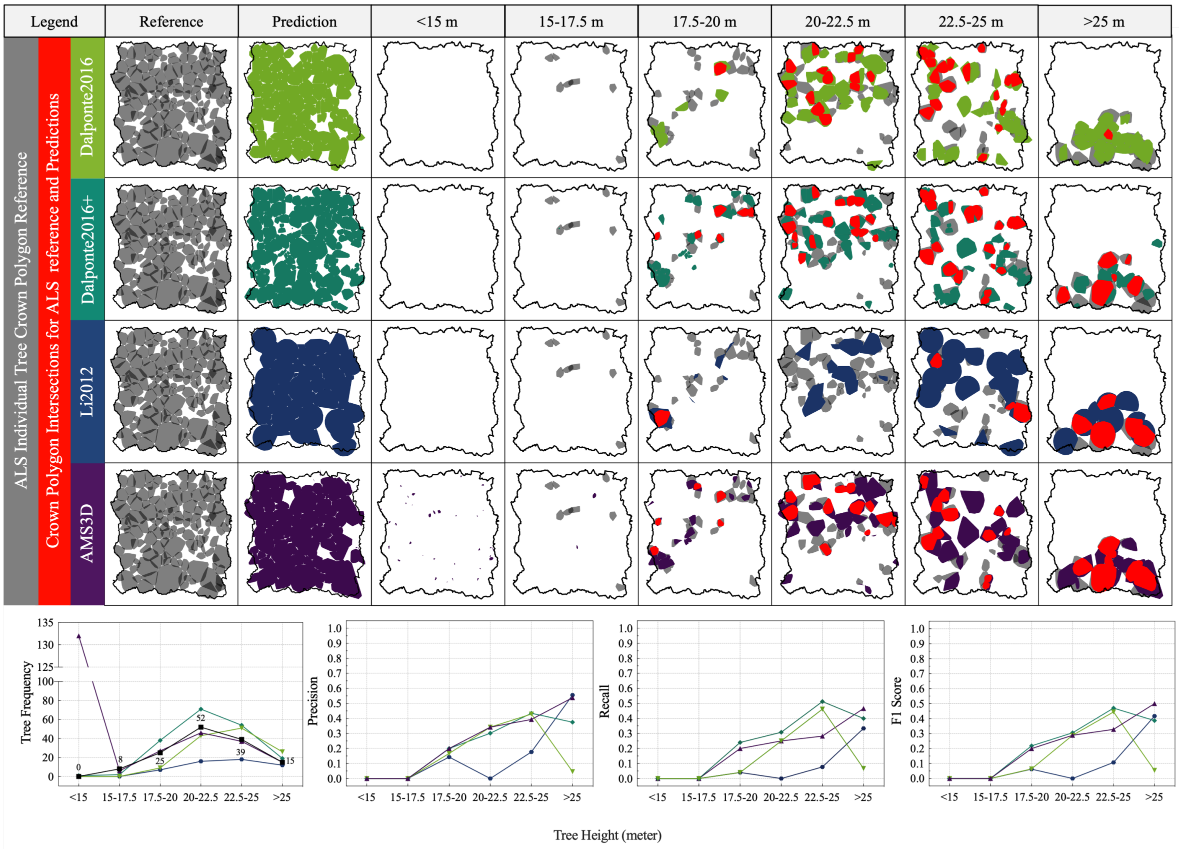


**Fig. H.2.** Precision, Recall and F1 Score changes with regards to tree height and confidence score (7 and 6) in Wytham Woods

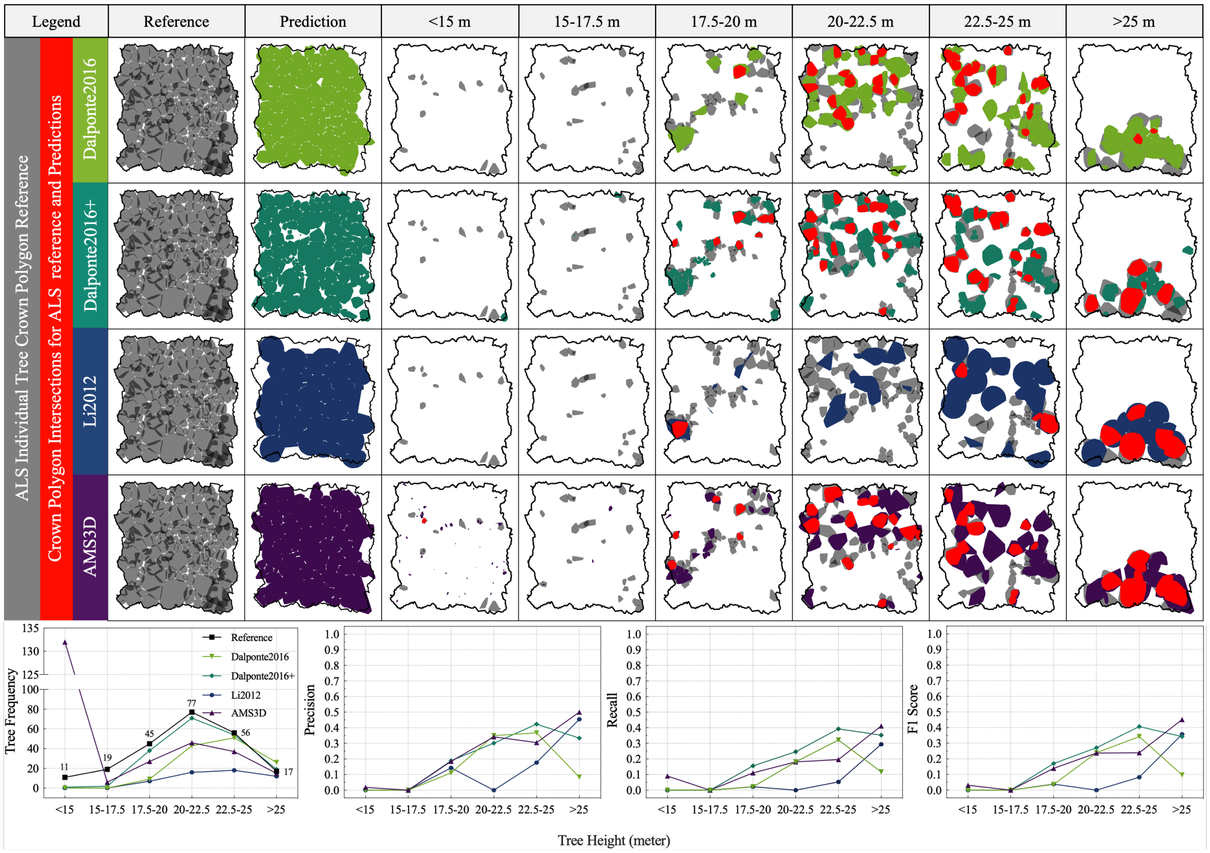


**Fig. H.3.** Precision, Recall and F1 Score changes with regards to tree height and confidence score=7,6,5 in Wytham Woods

### I Tau's contributions to Dalponte2016+'s performance

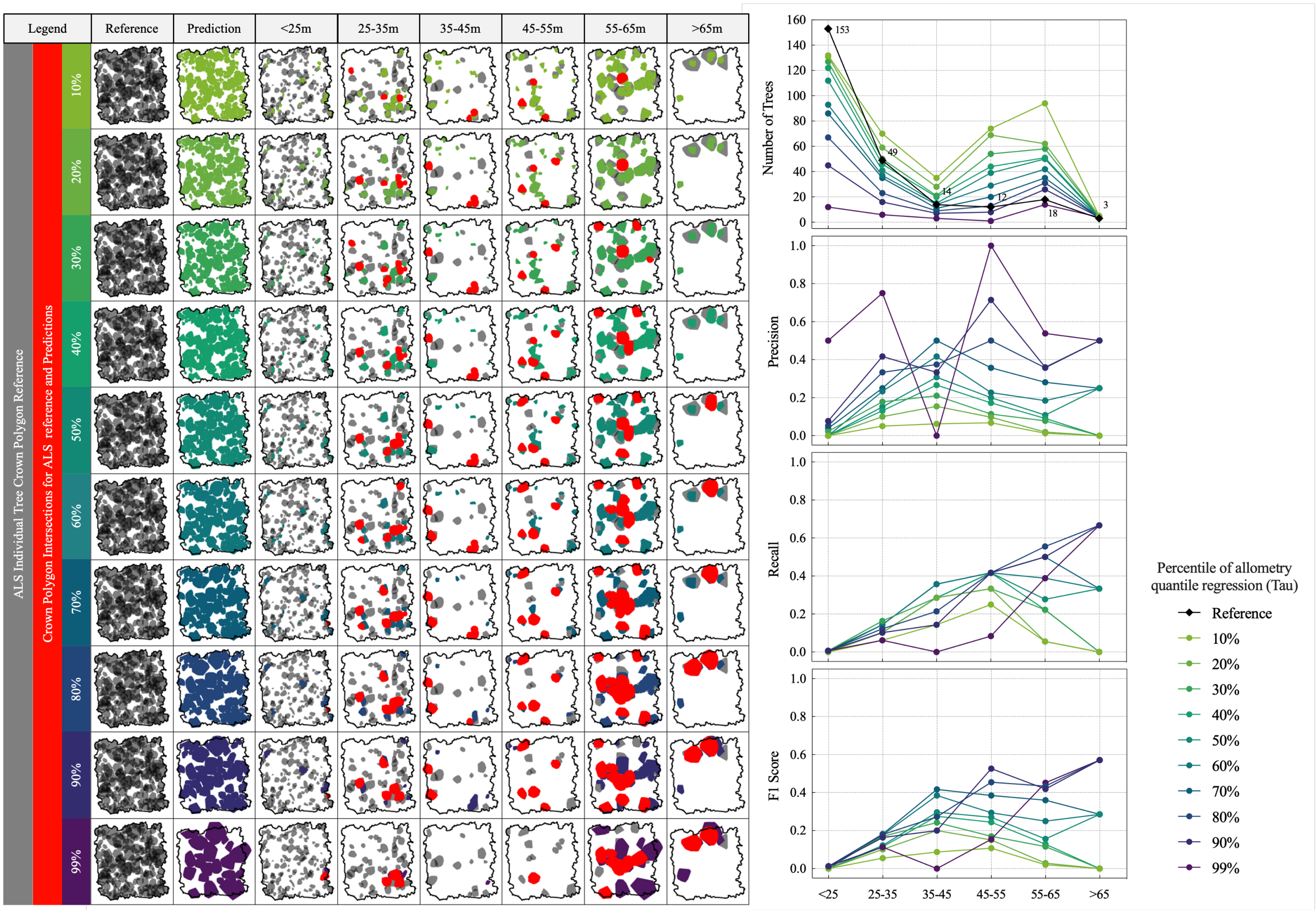


**Fig. I.1.** Dalponte2016+'s performance changes in Sepilok Forest with Taus and confidence score=7,6,5

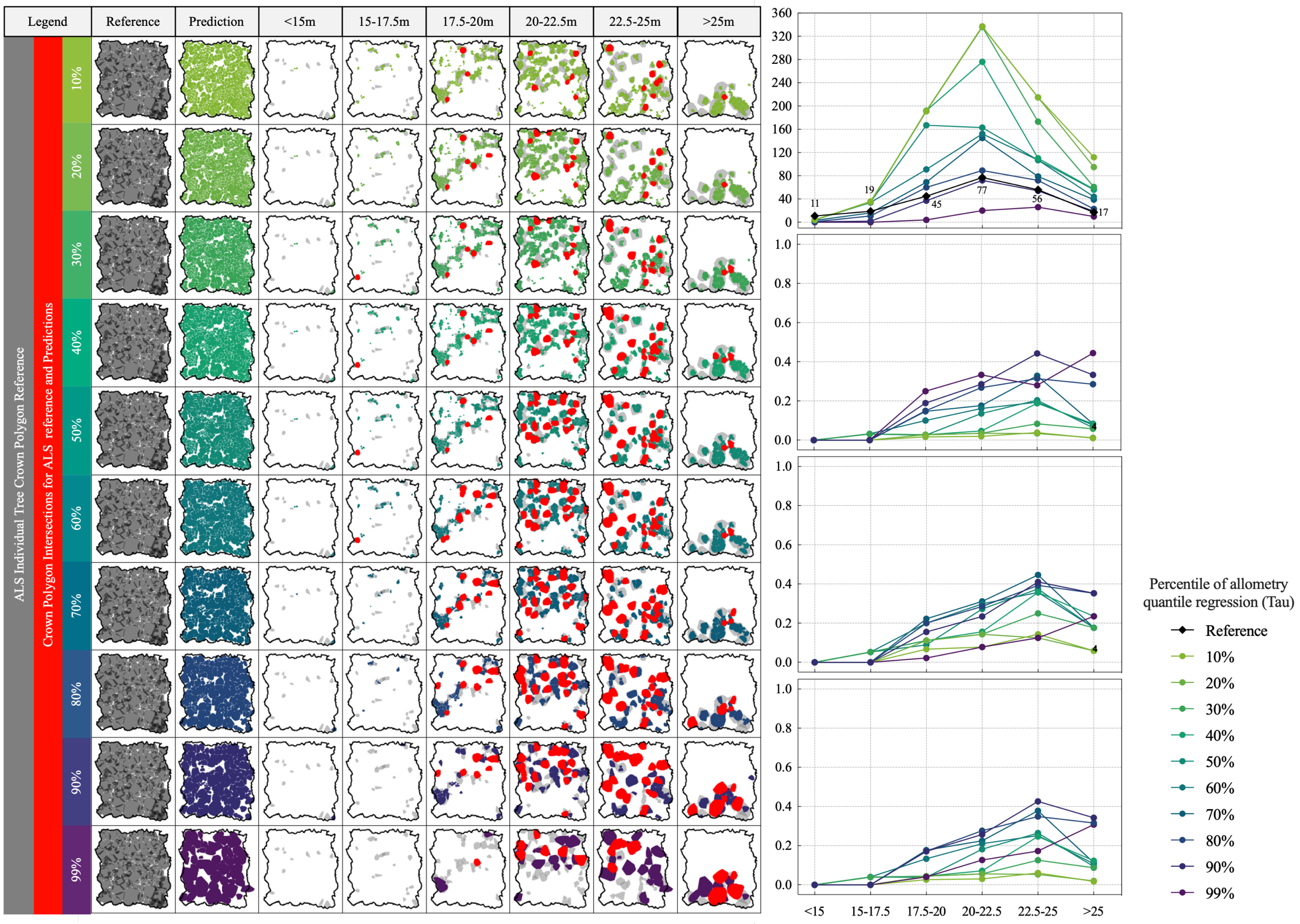


**Fig. I.2.** Dalponte2016+'s performance changes in Wytham Woods with Taus and confidence score=7,6,5

### J Crown diameter, tree height and IoU relations

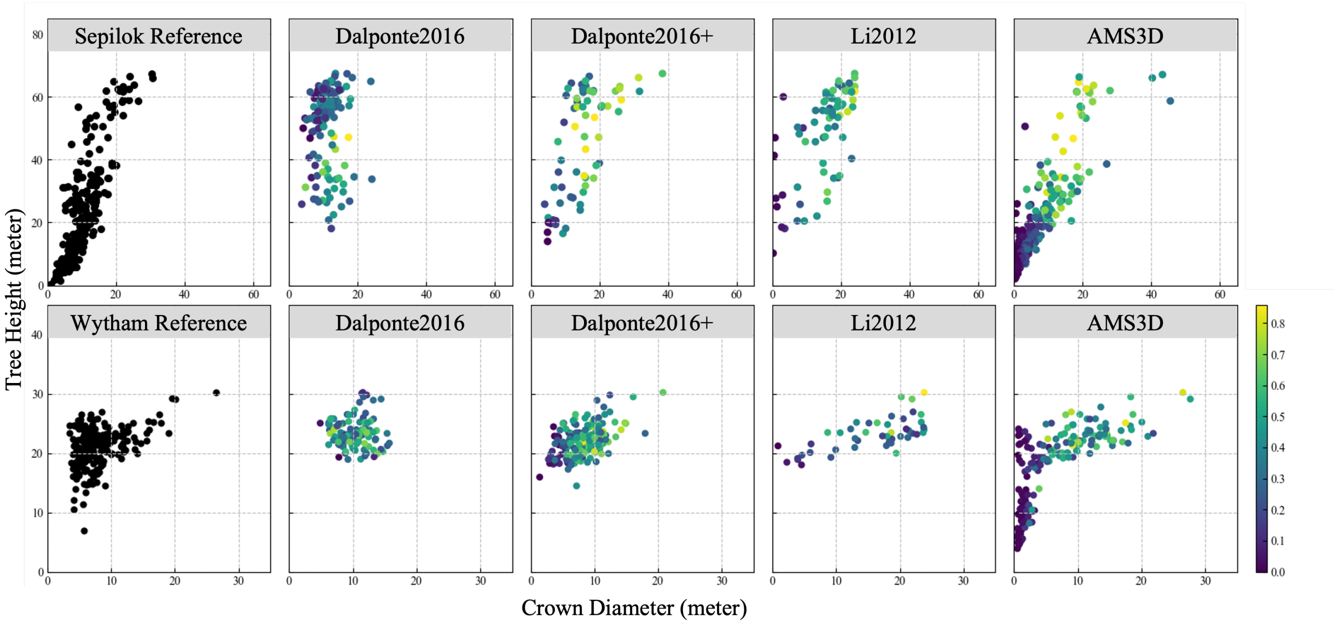


**Fig. J.1.** Allometry distribution scatters with regards to IoU; Scatters are colorized according to maximum IoU of predicted tree crown polygons with reference tree crown polygons; The first row represents Sepilok Forest, while the second row stands for Wytham Woods; Two plots in the first columns are reference allometry scatters, and other columns corresponds allometry scatters from different ITS algorithm with optimal parameter combination.
